# snRNA-seq of Huntington’s disease mice reveals vulnerability profiles of cortical cell types

**DOI:** 10.1101/2025.11.28.691138

**Authors:** Dennis Feigenbutz, Kerstin Voelkl, Rüdiger Klein, Irina Dudanova

**Affiliations:** Molecular Neurodegeneration Group, Max Planck Institute for Biological Intelligence, Martinsried, Germany; Department of Molecules – Signaling – Development, Max Planck Institute for Biological Intelligence, Martinsried, Germany; Center for Anatomy, Faculty of Medicine and University Hospital Cologne, University of Cologne, Cologne, Germany; Cologne Excellence Cluster on Cellular Stress Responses in Aging-Associated Diseases (CECAD), University of Cologne, Cologne, Germany

**Keywords:** Huntington’s disease, primary motor cortex, R6/2 mouse model, single-nucleus RNA-sequencing, ER-phagy

## Abstract

A common feature of neurodegeneration is selective vulnerability, where certain neurons succumb to disease, while others remain spared. The molecular underpinnings of these differences remain elusive. Here, we performed transcriptomic profiling of the motor cortex in a mouse model of Huntington’s disease (HD), an incurable hereditary movement disorder caused by a CAG repeat expansion in the Huntingtin gene. Strikingly, single-nucleus RNA-sequencing revealed a clear transcriptomic separation of HD and control samples within the vulnerable glutamatergic, but not disease-resistant GABAergic cell clusters. Tissue sampling at different time points allowed us to delineate a two-stage disease trajectory with distinct changes at early and late stages. Analysis of differentially expressed genes demonstrated progressive dysregulation of neuronal cell-type identity and upregulation of ER-phagy receptors. Mechanistic investigations in cellular HD models revealed increased ER-phagy, while knockdown of the ER-phagy receptor Tex264 resulted in elevated levels of the ER stress marker BiP, suggesting a protective role of ER-phagy in HD. Taken together, these findings advance our understanding of differential neuronal vulnerability, and identify ER-phagy as a new pathway in HD pathogenesis.

## Introduction

Huntington’s disease (HD) is a hereditary movement disorder with psychiatric and cognitive impairments (Tabrizi et al., 2020). The disease is currently incurable and inevitably lethal. The cause of HD is a pathological CAG triplet expansion in exon 1 of the Huntingtin (*HTT*) gene over the threshold of 36 repeats (The Huntington’s Disease Collaborative Research Group, 1993). This mutation results in an elongated polyglutamine (polyQ) tract in the HTT protein, leading to its aggregation and deposition of characteristic mHTT inclusion bodies in the brain. mHTT compromises multiple important cellular processes (Saudou and Humbert, 2016; Tabrizi et al., 2020), but the exact mechanisms leading to neuronal dysfunction and degeneration are not fully clarified. A thorough understanding of the cellular and molecular pathological events, especially those occurring at the early stages of disease progression, is essential for development of efficient therapies.

The most vulnerable cell types in HD are spiny projection neurons in the striatum and glutamatergic principal cells in the neocortex, although other cell types and brain regions also become affected as the disease progresses (Waldvogel et al., 2015). Studies in human gene mutation carriers and HD animal models have emphasized the role of cortical dysfunction and impaired corticostriatal communication at early stages of HD (Estrada-Sanchez et al., 2015; Estrada-Sanchez and Rebec, 2013; Gu et al., 2007; Reading et al., 2004; Schippling et al., 2009; Waldvogel et al., 2012). Motor cortical areas in particular have emerged as regions with a central role in pathogenesis (Fernandez-Garcia et al., 2020; Orth et al., 2010; Rosas et al., 2005; Rosas et al., 2008; Schippling et al., 2009).

Here, we have performed a time-resolved single-nucleus RNA-sequencing (snRNA-seq) profiling of the motor cortex in the R6/2 mouse model of HD at different disease stages from presymptomatic to advanced. We find a cell type-specific, biphasic pattern of transcriptional dysregulation, with cell types vulnerable to HD showing the most prominent alterations. Our analyses of differentially expressed genes and mechanistic follow-ups in cellular models furthermore point to ER-phagy as a new pathway with a potential neuroprotective role in HD.

## Results

### Glutamatergic neurons undergo a transcriptomic shift in HD mice

To investigate progressive transcriptional changes at different stages of HD, we used R6/2 mice, a transgenic fragment HD model expressing mHTT-exon1 under the human HTT promoter, characterized by an early onset and fast disease progression (Mangiarini et al., 1996). Primary motor cortex was harvested from transgenic mice and wildtype (WT) littermate controls at 3 different ages, representing different disease stages (Hosp et al., 2017): 5 weeks, a presymptomatic time point; 8 weeks, close to disease onset; and 12 weeks, when the mice are fully symptomatic (Fig. 1A). We isolated nuclei from the collected tissue and performed snRNA-seq using the droplet-based 10x Chromium system. After applying common quality control metrics (see Materials and Methods) and removing doublets, we retrieved a total of 95,074 nuclei across all samples (Fig. S1A). To annotate the nuclei, we integrated our dataset into the reference atlas of the mouse primary motor cortex from the BRAIN Initiative Cell Census Network (BICCN) (Yao et al., 2021). We chose to use the dataset snRNA-seq 10x v3 B, because it was also generated with the 10x technology and employed an improved nucleus isolation procedure. Prediction scores of nuclei being assigned to the correct cell type (e.g. L2/3 IT_1, L2/3 IT_2, etc.) (mean=0.84, sd=0.017 for WT, mean=0.8, sd= 0.041 for R6/2; Fig. S1B-C) and correct cell subclass (e.g. L2/3 IT, L5 IT, etc.) (mean=0.97, sd=0.004 for WT, mean=0.95, sd=0.018 for R6/2; Fig. S1C) were reliably high. Accordingly, visual inspection showed that nuclei of both genotypes from our dataset integrated well into the clusters of the reference atlas (Fig. S1D), with the exception of Layer 6b (L6b) glutamatergic neurons that were scarce in our dataset. This might be due to differences in microdissection, as L6b is the deepest cortical layer.

**Figure 1.**
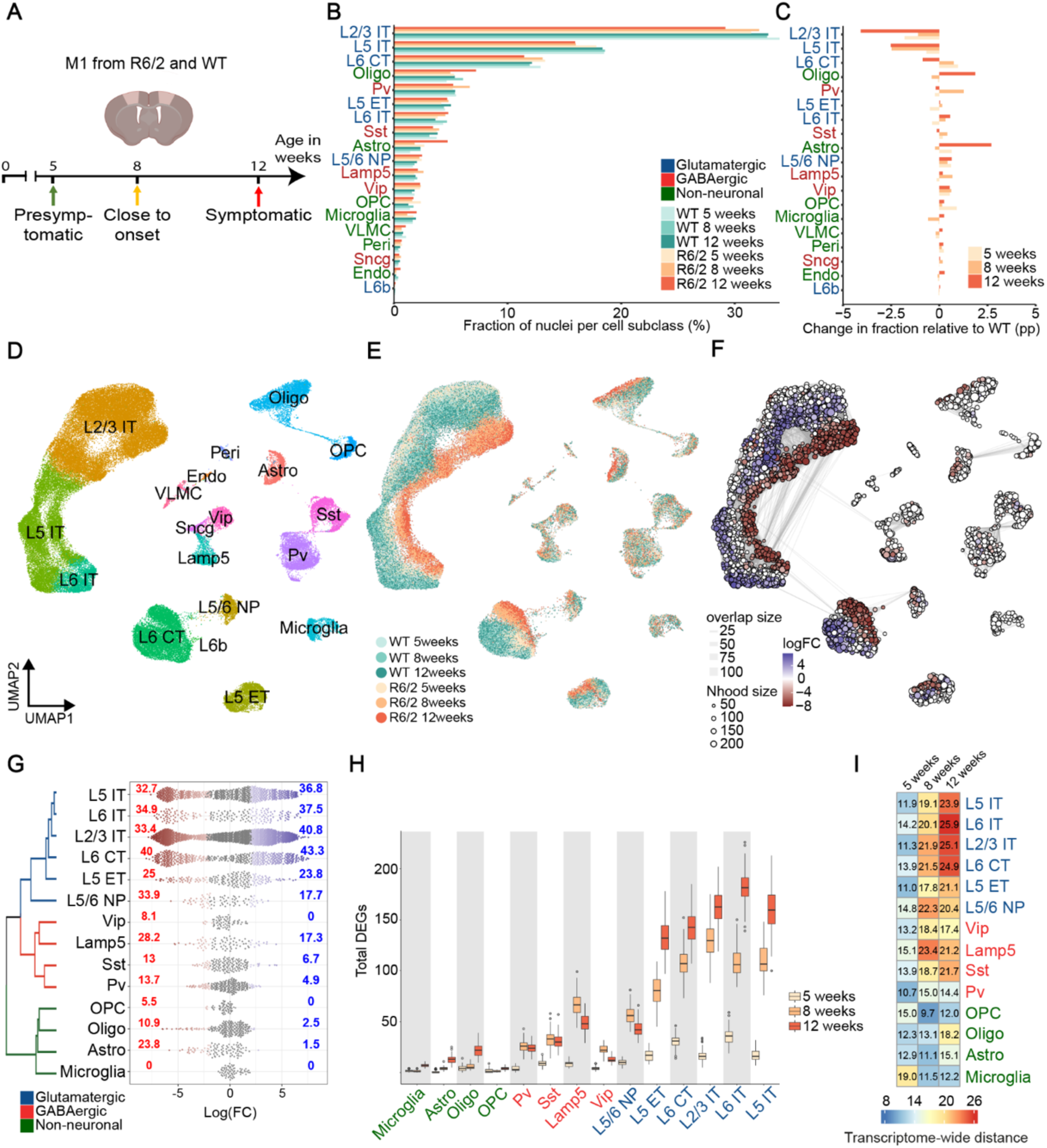
snRNAseq of the primary motor cortex of R6/2 mice reveals transcriptomic shift of glutamatergic neurons. **A**, Experimental design. **B**, Fraction of nuclei of each cell subclass at different time points. **C**, Change in fraction of R6/2 nuclei per cell subclass relative to the respective WT control in percentage points (pp). **D**, UMAP visualization of nuclei clusters including R6/2 and WT nuclei from all time points, color-coded by cell subclass. **E**, UMAP visualization of nuclei clusters color-coded by time point and genotype. Note the transcriptomic shift in glutamatergic neuron subclasses. **F**, UMAP plot with nuclei grouped in neighborhoods. Neighborhoods significantly enriched for WT nuclei are shown in blue and for R6/2 nuclei in red. **G**, Swarm plot of neighborhoods per cell subclass. For differentially abundant neighborhoods (alpha=0.1), the Log(FC) enrichment for WT (blue) or R6/2 (red) nuclei is indicated. Neighborhoods without significant over-representation are shown in grey. **H**, Box plots of iterative differential gene expression analysis after subsampling 300 nuclei per cell subclass for different time points. **I**, Quantification of transcriptome-wide distance (“disease score”) between R6/2 and WT nuclei of indicated cell subclasses.

Based on this annotation, we identified 12 neuronal and 7 non-neuronal cell subclasses in our dataset (Fig. 1B, D). Among the neuronal subclasses, 7 were glutamatergic and 5 GABAergic. Glutamatergic clusters included L2/3, L5 and L6 intratelencephalic (L2/3 IT, L5 IT and L6 IT) neurons, L5 extratelencephalic (L5 ET) neurons, L6 corticothalamic (L5 CT) neurons, L6b neurons, and L5/6 near-projecting (L5/6 NP) neurons. GABAergic clusters were represented by Parvalbumin (Pv), Somatostatin (Sst), vasoactive intestinal peptide (Vip), Lysosomal associated membrane protein family member 5 (Lamp5) and Synuclein Gamma (Sncg) interneurons. Non-neuronal subclasses were comprised of oligodendrocytes, oligodendrocyte precursor cells (OPCs), astrocytes, microglia, endothelial cells, pericytes and vascular leptomeningeal cells. Cell subclasses with fewer than 200 nuclei per time point (L6b glutamatergic neurons, Sncg GABAergic neurons, endothelial cells, pericytes and vascular leptomeningeal cells) were excluded from many of the further analyses due to insufficient statistical power.

We observed a high correlation among WT transcriptomes of different cell subclasses within the glutamatergic class and GABAergic class, respectively. Glial cell subclasses also exhibited positive correlation values between each other, but negative correlation values with neuronal subclasses (Fig. S2A). All cell subclasses identified in our dataset showed the typical marker expression described in previous scRNA-seq datasets of the primary motor cortex (Fig. S2B-D) (Yao et al., 2021; Yao et al., 2023). For example, the gene solute carrier family 17 member 7 (*Scl17a7*) was highly expressed across all glutamatergic neuron subclasses (Fig. S2B, D), whereas glutamic adic decarboxylase 2 (*Gad2*) was specific to GABAergic neuron subclasses (Fig. S2C-D).

We next examined the cellular composition of all samples. The fractions of most cell subclasses were similar in samples of different genotype and age, with the exception of IT glutamatergic neurons, oligodendroglia and astrocytes. The L5 IT cell subclass showed a ∼15% reduction in R6/2 mice already at 8 weeks and the L2/3 IT subclass at 12 weeks. In contrast, oligodendroglia numbers were 1.5-fold higher and astrocyte numbers two-fold higher in 12-week-old R6/2 mice (Fig. 1B-C).

UMAP visualization of the dataset revealed a striking transcriptomic shift of the nuclei of 8-week and 12-week HD mice compared to 5-week old HD mice and WT littermates. Remarkably, the shift occurred in all glutamatergic neuronal cell subclasses, but not in any of the GABAergic cell subclasses (Fig. 1E). Among the glutamatergic neurons, the strongest effect was seen in IT neurons, which project to the striatum and are severely affected in HD (McColgan et al., 2020), as well as in L6 corticothalamic (CT) neurons. Consistently, differential abundance testing (Dann et al., 2022) showed that in most glutamatergic neuronal cell subclasses, there was an increased number of neighborhoods enriched for either R6/2 or WT nuclei, while in GABAergic and non-neuronal cell subclasses the neighborhoods mostly contained nuclei of mixed genotypes (Fig. 1F). Glutamatergic IT cell subclasses exhibited especially high over-representation values of single genotypes (Fig. 1F-G). These results point to a different extent of transcriptional dysregulation in different cortical cell types, with glutamatergic neurons and IT neurons in particular undergoing the greatest disease-related changes.

To further characterize the transcriptomic changes in cortical cell subclasses and determine their temporal progression, we employed two quantification methods. In the first approach, we performed 100 iterations of randomly subsampling 300 nuclei from each cell subclass to compensate for the varying numbers of nuclei for individual cell subclasses, and quantified the numbers of differentially expressed genes (DEGs). Neurons showed overall more DEGs than non-neuronal cells, glutamatergic neurons more than GABAergic ones, and among the glutamatergic neurons all the IT cell subclasses displayed the highest numbers of DEGs (Fig. 1H). DEG numbers increased with age for most cell subclasses, with the greatest change occurring between 5 and 8 weeks of age.

In a second approach, we calculated the transcriptome-wide distance between WT and R6/2 nuclei across different subclasses at each time point (“disease score”) (Pineda et al., 2021). We observed a progressive, age-dependent increase in disease score in neuronal cell subclasses, with especially high values in glutamatergic IT cell subclasses and in L6 CT neurons (Fig. 1I). Taken together, these results demonstrate that the extent of transcriptomic changes increases with disease progression and markedly differs between cortical cell populations, in line with differential cellular vulnerability to HD.

### Biphasic progression of transcriptional changes in HD cortex

For further analyses, we focused on glutamatergic IT neuron types, as they displayed the most pronounced transcriptomic alterations in HD mice. To investigate age-dependent disease effects, we applied pseudotime trajectories to the nuclei using the Slingshot package (Street et al., 2018). WT nuclei of IT neurons of different ages were evenly distributed along the pseudotime trajectory, while R6/2 nuclei were markedly shifted. 12-week R6/2 nuclei occupied the end of the trajectory with the highest pseudotime values, while 8-week R6/2 nuclei were found in an intermediate position between WT and 12-week R6/2. Surprisingly, 5-week R6/2 nuclei were shifted in the opposite direction, showing lower pseudotime values than WT nuclei (Fig. 2A-D). This biphasic pattern was observed for nuclei of all IT neuron subclasses as well as other glutamatergic subclasses (Fig. S3A-C). In line with these findings, the correlation of transcriptomic changes in most glutamatergic cell subclasses was low between 5 and 8 week samples, but high between 8 and 12 week samples (Fig. 2G-H). In contrast to glutamatergic neurons, GABAergic neurons and glial cells of different ages showed overlapping distribution in UMAP space, so that no disease-related trajectory could be mapped (Fig. S3D). Taken together, these data point to a biphasic progression of disease signatures in glutamatergic neurons, with distinct transcriptomic alterations at early and late stages.

**Figure 2.**
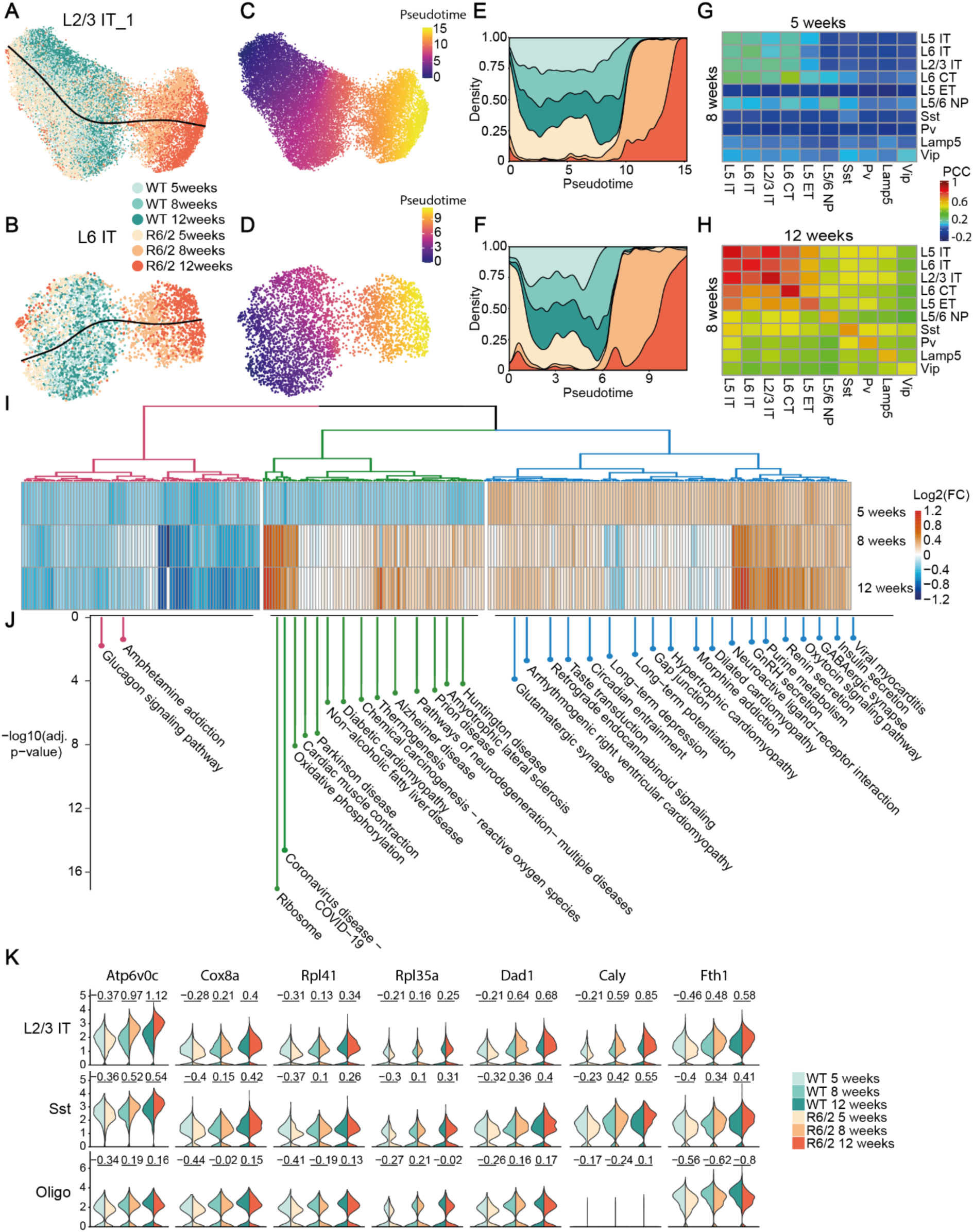
Biphasic temporal progression of HD-related transcriptomic changes. **A-B**, Nuclei of the L2/3 IT_1 cell type (A) and L6 IT cell subclass (B) color-coded for age and genotype were plotted with slingshot trajectory. **C-D**, Nuclei of the L2/3 IT_1 cell type (C) and L6IT cell subclass (D) color-coded by their pseudotime value. **E-F**, Density plot of pseudotime values of the L2/3 IT_1 nuclei (E) and L6 IT nuclei (F). **G**, Heat map of pairwise Pearson correlation of transcriptomic change in R6/2 relative to WT mice for all genes per cell subclass between 8 and 5 weeks. **H**, Heat map of Pearson correlation of transcriptomic change for all genes per cell subclass between 8 and 12 weeks. **I**, Heat map of gene expression in L2/3 IT at different ages. Genes were hierarchically clustered by similarity of temporal expression and separated into clusters by k-means clustering (k=3). **J**, KEGG pathways analysis for each cluster from **I**. **K**, Violin plots of normalized expression levels for example genes that are downregulated at 5 weeks and upregulated at 8 and 12 weeks in glutamatergic L2/3 IT neurons. For comparison, plots of the same genes in Sst neurons and oligodendrocytes are shown. The biphasic pattern is less pronounced in Sst cells and absent in oligodendrocytes. Values above the violin plot indicate Log2(FC).

We next asked which genes drive these biphasic changes during disease progression. Hierarchical clustering of all DEGs identified at the 5 weeks time point by their expression changes in R6/2 mice at 5, 8 and 12 weeks revealed three sets of genes. Two sets contained genes whose differential expression at 5 weeks positively correlated with their differential expression at 8 and 12 weeks, being either consistently downregulated (first cluster) or upregulated (third cluster) at all ages (Fig. 2I). Among consistently downregulated genes, we found such categories as “Glucagon signaling pathway” and “Amphetamin addiction”. Among consistently upregulated genes, Kyoto Encyclopedia of Genes and Genomes (KEGG) pathway analysis identified categories such as “Glutamatergic synapse”, “Long-term depression” and “Long-term potentiation”. (Fig. 2J). In contrast, the remaining set of genes (second cluster in Fig. 2I) showed opposite expression changes at early and late stages, being downregulated at 5 weeks and upregulated at 8 and 12 weeks. According to KEGG pathway analysis, this gene set included such gene categories as “Ribosome” and “Oxidative phosphorylation”, as well as several categories involved in neurodegenerative diseases including Parkinson’s disease, Alzheimer’s disease, prion diseases and HD (Fig. 2J). Examples of genes exhibiting clear bidirection regulation in glutamatergic cell types include ATPase H+ transporting V0 subunit c (*Atp6v0c*), cytochrome c oxidase subunit 8A (*Cox8a*), ribosomal proteins L41 (*Rpl41*) and L35a (*Rpl35a*), Calcyon neuronspecific vesicular protein (*Caly*) and Ferritin heavy chain 1 (*Fth1*) (Fig. 2K). These results demonstrate biphasic progression of transcriptional changes in the HD cortex, with distinct sets of regulated genes at early and late stages of the disease.

### Differential gene expression in neuronal cell subclasses

In contrast to the 5-week time point, the transcriptomic changes found in 8- and 12-week-old samples were largely overlapping. 49% upregulated DEGs and 42% downregulated DEGs were common between these two time points, while the total numbers of DEGs increased with age (Fig. 3A, B, Supplementary Tables S1-S3). To reveal common and cell type-specific disease signatures, we therefore focused on the 12-week time point. Interestingly, most upregulated DEGs were shared between different subclasses of glutamatergic neurons (Fig. 3C). In contrast, downregulated DEGs were largely cell subclass-specific (Fig. 3D). Thus, 391 upregulated (27%) and only 96 (10%) downregulated DEGs were common to at least 5 out of 6 glutamatergic neuron subclasses. To explore the expression of subclass-specific genes in more detail, we identified the top 100 marker genes for each cell subclass and monitored their expression across the disease stages. Strikingly, we found a pronounced decrease in the expression of marker genes in all cell subclasses except microglia and OPCs between 5 and 8 weeks, and for many of them a further decline between 8 and 12 weeks (Fig. 3E). We then examined the expression of single molecular markers commonly used to identify cortical neuronal subclasses. The characteristic markers of the major GABAergic interneuron subclasses *Pv*, *Sst*, *Vip*, and *Lamp5* showed a clear downregulation, especially at the 12-week time point. Likewise, the glutamatergic subclass markers Neurogranin (*Nrgn*, L2/3 IT), neural EGFL like 1 (*Nell1*, L5 IT), RNA binding fox-1 homolog 1 (*Rbfox1*, L6 CT) and TAFA chemokine-like family member 1 (*Tafa1*, ET) were downregulated in R6/2 mice (Supplementary Fig. S4A).

**Figure 3.**
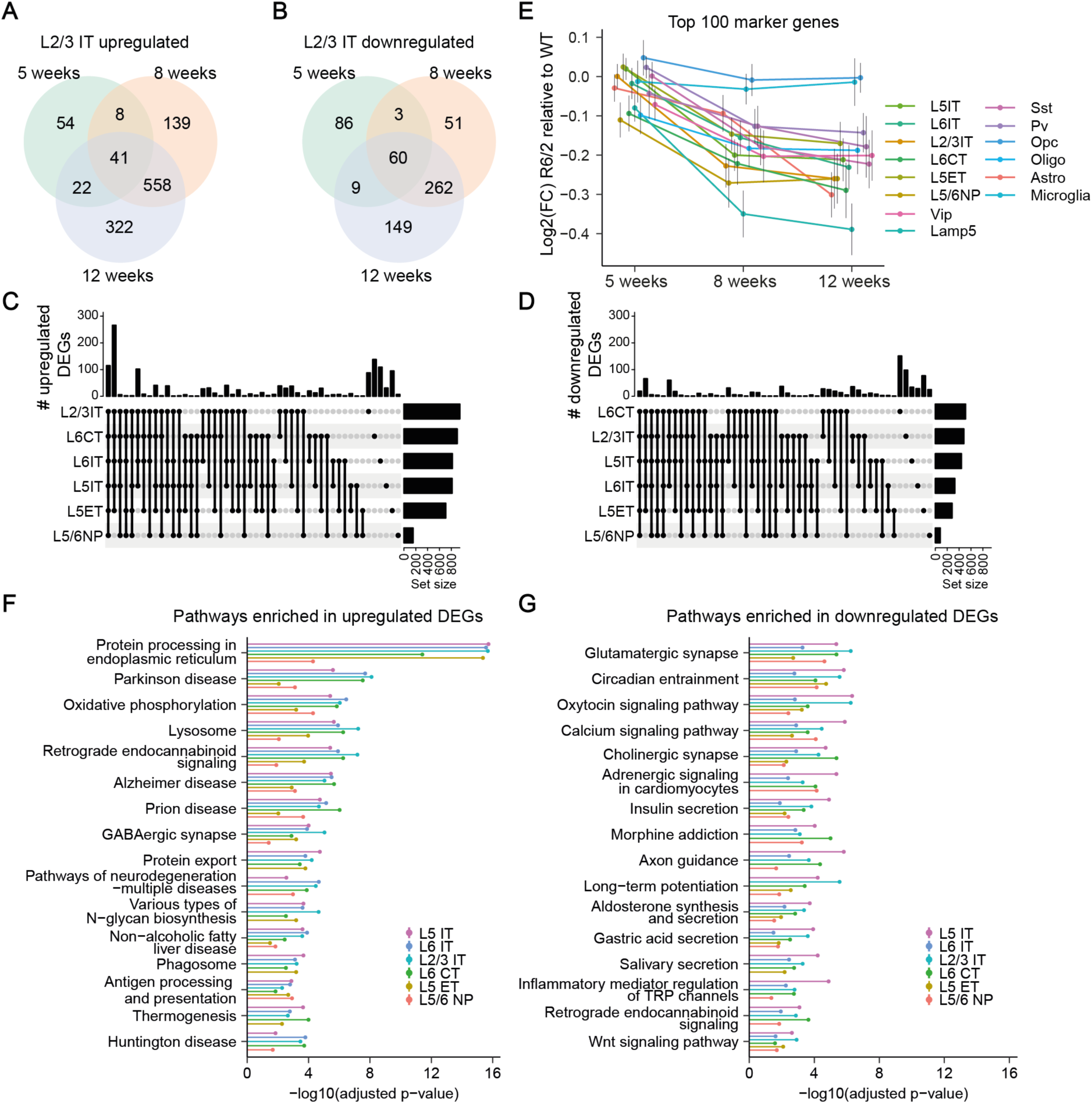
Analysis of DEGs and their pathways. **A-B**, Venn diagrams of upregulated (A) and downregulated (B) DEGs in L2/3 IT neurons. **C-D**, UpSet plots of upregulated (C) and downregulated (D) DEGs at 12 weeks for glutamatergic neuron subclasses. **E**, Average dysregulation for the top 100 marker genes for each cell subclass in R6/2 mice at different ages. Error bars represent 95% confidence interval. **F-G**, KEGG pathway analysis of upregulated (F) and downregulated (G) DEGs for gutamatergic neuron subclasses at 12 weeks. Pathways are ranked by the average adjusted p-value for all excitatory cell types and the top 15 pathways are shown.

We hypothesized that the widespread decline in cell subclass marker genes could be due to changes in the expression of key transcription factors that define neuronal cell type identity. We focused on the recently described set of transcription factors that regulate the identity of glutamatergic and GABAergic neuronal subclasses in the telencephalon (pallium- and subpallium-specific transcription factors, respectively) (Yao et al., 2023) and analyzed their expression in the HD cortex. Interestingly, we observed a decrease in the levels of pallium-specific transcription factors in all glutamatergic subclasses (Supplementary Fig. S4B). Among the transcription factors that were particularly strongly downregulated, we found *Neurod1*, FEZ family zinc finger 2 (*Fezf2*), JunB proto-oncogene (*Junb*), FosB proto-oncogene (*Fosb*), zinc finger E-box binding homeobox 2 (*Zeb2*), inhibitor of DNA binding 2 (*Id2*) and nuclear factor 1 X (*Nfix2*) (Supplementary Fig. S4C). In contrast to glutamatergic cell subclasses, the expression of subclass-defining transcription factors in GABAergic neuron subclasses was mostly unchanged (Supplementary Fig. S4B-C). These findings suggest a general defect in maintaining subclass-specific gene expression in the HD cortex, which in glutamatergic neurons was accompanied by downregulation of subclass-specific transcription factors.

### Pathway analysis implicates ER protein homeostasis in HD pathogenesis

To uncover specific molecular pathways relevant for HD pathogenesis, we next performed KEGG pathway analysis of DEGs identified in R6/2 mice at 12 weeks of age. For the upregulated genes, the most significant KEGG pathway category was “Protein processing in the endoplasmic reticulum”, followed by “Oxydative phosphorylation” and other pathways related to protein processing (“Lysosome”, “Protein export”) and to neurodegenerative proteinopathies (“Parkinson’s disease”, “Alzheimer’s disease”, “Prion disease”) (Fig. 3F). In agreement with “Protein processing in the endoplasmic reticulum” being the most significantly dysregulated pathway, we found a variety of endoplasmic reticulum (ER)-related genes to be significantly upregulated, including ER chaperones (*Hspa5, Canx, Calr*), glycosylation enzymes (*Dad1, Tusc3, Rpn1*), ER translocation proteins (*Sec62*), ER-associated degradation (ERAD) components (*Erlec1, Ubxn4, Os9*) and receptors for ER-specific autophagy (ER-phagy) (*Tex264, Rtn3, Sec62*). In addition to the ER protein quality control machinery, several other chaperones (*Hsp90ab1, Hsp90aa1, Hspa8, Pcsk1n*) and lysosomal genes (*Lamp1, Lamp2, Ctsd*) were also among the upregulated DEGs, highlighting the role of the protein homeostasis network in HD pathogenesis (Fig. S5A-D, Table 3). In contrast, the KEGG pathways for downregulated genes were often related to neuron-specific functions, including “Glutamatergic synapse”, “Cholinergic synapse”, “Axon guidance” and “Long-term potentiation” (Fig. 3G). Accordingly, among the downregulated DEGs we found many genes encoding structural components of synapses (*Homer1*, *Camk2a, Camk4, Grin2a*) and axon guidance molecules (*Unc5d, Sema4a*) (Fig. S5B-C, Table 1). In summary, upregulated genes are largely shared between neuronal subclasses and often related to ER protein quality control, while downregulated genes are more cell subclass-specific and often involved in neuron-specific functions.

### A gene module with putative neuroprotective genes

To identify disease-related gene networks and key genes that are central to these networks (hub genes), we applied weighted gene co-expression network analysis (WGCNA), which groups genes into modules based on the similarity of their expression across genotypes and time points (Feregrino and Tschopp, 2022). We identified 20 co-expression modules containing between 65 and 431 genes (Fig. 4A). Correlation of the module eigengene expression with the genotype at each time point revealed a number of positively correlated modules, whose genes were mostly upregulated in R6/2 mice, and several negatively correlated modules, whose genes were largely downregulated in R6/2 mice (Fig. 4B). Consistent with the similarity of transcriptomic changes observed at 8 and 12 weeks (Fig. 2H, I), correlation values of the modules were also similar at these two time points. The turquoise module, which exhibited a negative correlation at 5 weeks and a positive correlation at 8 and 12 weeks (Fig. 4B), contained genes with biphasic changes in transcription during disease progression (Fig. 2I-K), for example *Atp6v0c*, *Cox8a*, *Rpl35a*, *Dad1*, *Caly*, and ubiquitin B *(Ubb)*.

**Figure 4.**
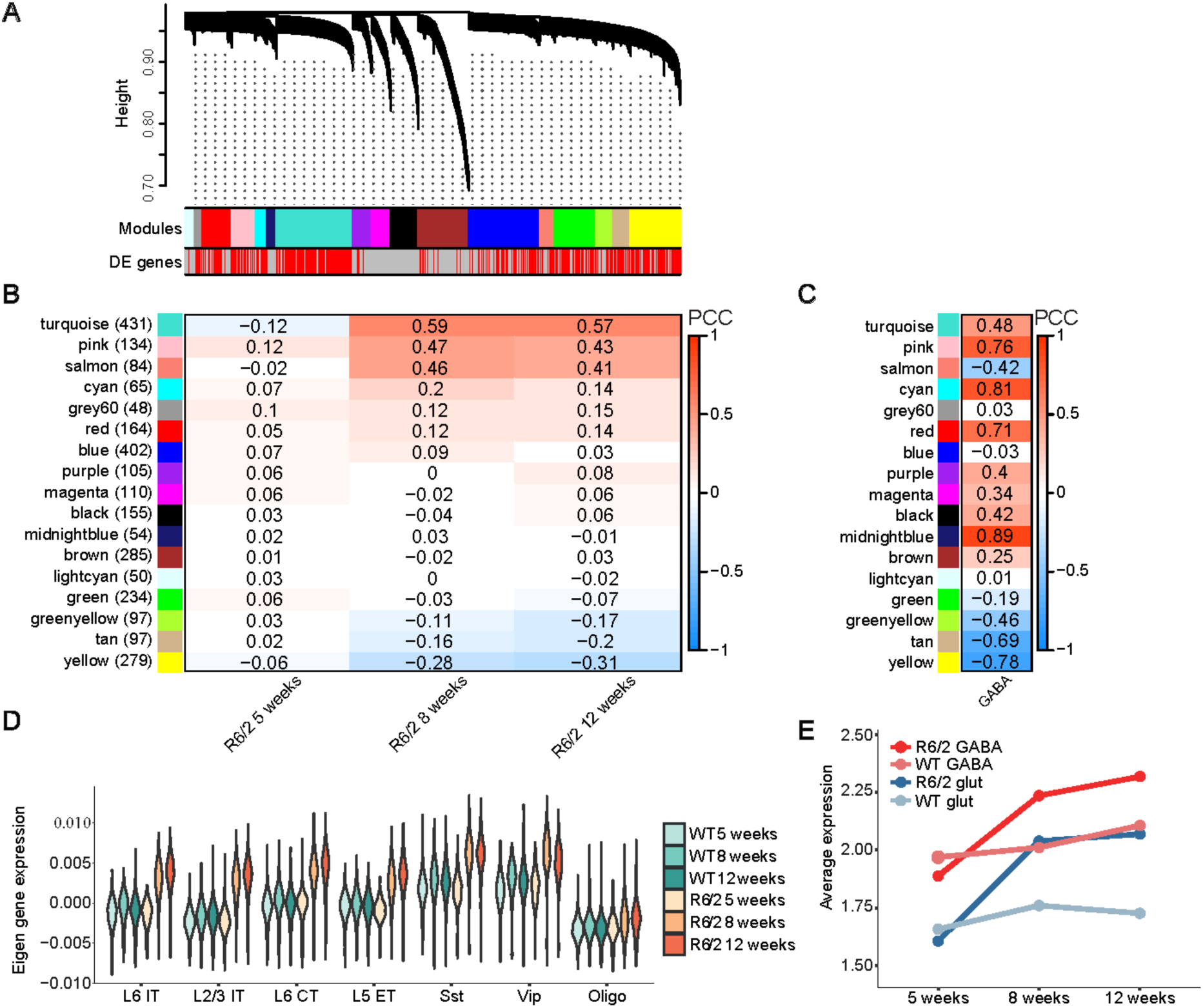
scWGCNA identifies modules of co-expressed genes and their hub genes. **A**, Gene dendrogram showing the co-expression modules identified with scWGCNA. **B**, Pearson correlation of the module eigengenes with the R6/2 genotype across different ages. **C,** Pearson’s correlation of the module eigengenes with GABAergic WT nuclei. **D,** Module eigengene expression of the turquoise module in different cell subclass. **E**, Average expression of all genes of the turquoise module in WT and R6/2 GABAergic and glutamatergic neurons with disease progression.

We then examined the cell subclass specificity of the modules in healthy WT mice by correlating the expression of the module eigengenes to glutamatergic and GABAergic neuron subclasses (Fig. 4C). In this analysis, higher correlation values indicated higher expression levels of the modules in WT GABAergic neurons, while lower values corresponded to higher expression in WT glutamatergic neurons. Interestingly, many modules whose genes were upregulated in R6/2 mice (e.g. turquoise, pink, cyan, red) showed high correlation values, whereas modules downregulated in R6/2 mice (green, greenyellow, tan, yellow) had low correlation values (Fig. 4C). These data suggest that modules with higher basal expression levels in GABAergic neurons are the ones that become upregulated in HD. To expore this effect in more detail, we focused on the turquoise module, which showed the strongest correlation with the R6/2 genotype, and analyzed the average expression of its genes in different cell subclasses. In WT cortex, the expression of the turquoise module was consistently higher in GABAergic cell subclasses than in glutamatergic ones, without any age-related changes. In R6/2 animals, the expression levels in all subclasses were similar to controls at 5 weeks, but increased at 8 weeks, with the values of the glutamatergic cell subclasses of R6/2 mice becoming similar to the GABAergic cells of wiltype mice (Fig. 4D-E).

These observations raised the possibility that the turquoise module might contain genes that are neuroprotective in the context of HD. Their higher baseline expression levels in cortical GABAergic interneurons might contribute to the resilience of interneurons to HD, and the upregulation of these genes in R6/2 mice might represent a compensatory protective reaction to the emergent pathology. We therefore sought to test whether the turquoise module genes were among the genes essential for neuronal survival in a healthy brain and in HD conditions. To this end, we explored a dataset from an *in vivo* genome-wide screen for genes that are essential for neuronal survival in the WT mouse brain and genes that modify mHTT toxicity in HD models (Wertz et al., 2020). Out of the 3,875 neuronal essential genes identified in that study, 3,009 were detected in our dataset. Notably, the turquoise module showed a significant overrepresentation of neuronal essential genes, while mHTT toxicity modifiers were not significantly overrepresented in any of the modules (Supplementary Fig. S6A-B). Taken together, gene co-expression analysis releaved a module that was enriched for neuronal essential genes, had higher baseline expression in the disease-resistant GABAergic neurons and became upregulated in HD mice around disease onset. We propose that this module might contain genes that are neuroprotective in the context of mHTT toxicity.

To identify specific key factors with putative neuroprotective properties, we constructed a gene network containing the top 30% of hub genes from the modules that exhibited an absolute correlation value higher than 0.15 in at least one time point (Fig. 4B). Within the network, we plotted the 30% highest absolute adjacencies, so that tightly co-expressed genes appear closer to each other. In addition, we highlighted all neuronal essential genes and mHTT toxicity modifiers (Wertz et al., 2020). Among the upregulated genes (from the turquoise, pink salmon and cyan modules), we again found a number of chaperones, including ER chaperones (*Hspa5*, *Hsp90aa1*, *Hsp90ab1*, *Hsp90b1*) to have a central position in the network (Fig. 5). *Hspa5*, also known as BiP or GRP78, is the key ER-resident Hsp70 chaperone that, in addition it central role in protein folding, is pivotal for activating the unfolded protein response (UPR). The chaperone *Hspa8* also appeared as a hub gene, being at the same time a neuronal essential gene and a mHTT toxicity modifier. Among the downregulated genes (from the yellow, tan and greenyellow modules), we detected many genes related to synaptic function, such as *Camk2a*, glutamate receptor *Grin2a*, ryanodine receptor 2 (*Ryr2*), K_V_ voltage-gated channel interacting protein 4 (*Kcnip4*) and Ca_V_ voltage-gated channel auxillary subunit (*Cacna2d1*). These results are in line with the pathway analysis and highlight the key role of protein homeostasis- and synapse-related genes in HD.

**Figure 5.**
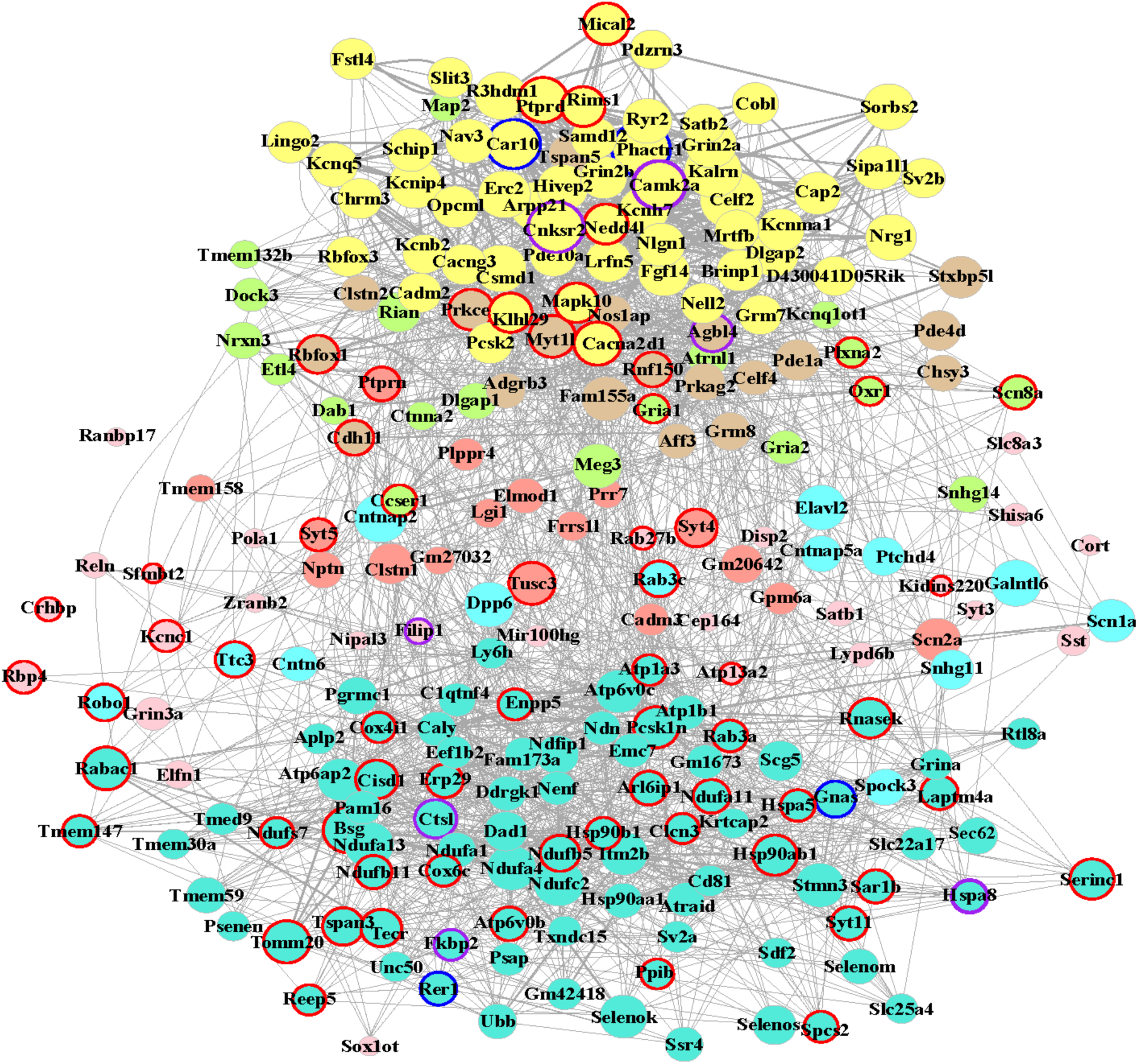
Gene co-expression network with hub genes of the R6/2 cortex transcriptome. Force-directed network representation of the top 30% hub genes of all modules with a correlation of more than 0.15 or less than -0.15 with the R6/2 genotype. The top 5% highest and lowest adjacency values were considered for network generation. Vertices are colored by the module the gene belongs to (see Fig. 4). Yellow, tan and green genes are part of downregulated modules; turquoise, pink salmon and cyan genes are part of upregulated modules. The size of the vertice is proportional to the centrality of the gene, e.i. larger vetices correspond to genes more tightly co-expressed with the other genes of the respective module. The length of the edges represents the adjacency of the genes, e.i. the closer two genes are, the more tightly they are co-expressed. Neuronal essential genes are indicated by red circles, mHTT toxicity modifiers by blue circles, and genes that belong to both categories by purple circles.

### mHTT causes an increase in ER-phagy

Our pathway and hub gene analyses revealed that ER protein homeostasis-related genes were strongly upregulated in the HD mouse cortex. Among those were a number of genes related to ER-phagy - the process of selective clearance of parts of the ER through the autophagy-lysosomal pathway. This process is important for maintaining ER homeostasis, particulary in conditions of ER stress and proteotoxicity (Reggiori and Molinari, 2022). However, the role of ER-phagy in HD has not yet been investigated. We therefore focused our further analyses on ER-phagy. This process requires ER-phagy receptors, which are ER-associated proteins that interact with the autophagy machinery. To date, six membrane-anchored ER-phagy receptors have been described in mammals: Tex264, Rtn3, Sec62, Ccpg1, Retreg1 and Atl3 (Reggiori and Molinari, 2022). Remarkably, five of these receptor transcripts (*Tex264, Rtn3, Sec62, Ccpg1 and Retreg1*) were significantly upregulated in R6/2 mice starting from 8 weeks of age, and *Rtn3* expression was already elevated at 5 weeks (Fig. 6A). In addition, further genes implicated in ER-phagy were also upregulated, such as *Pgrmc1* and *Calnexin* (*Canx*) (Tables 2-3). PGRMC1 is a binding partner of RTN3 that captures ER-phagy cargo proteins (Chen et al., 2021), and Calnexin is an ER chaperone that was shown to regulate ER-phagy in neurons (Wolf et al., 2024). Of note, three of the ER-phagy receptor genes, *Sec62*, *Tex264* and *Rtn3*, were part of the turquoise gene module with putative neuroprotective properties, and *Sec62* was also identified as a hub gene in the co-expression network (Fig. 5).

**Figure 6.**
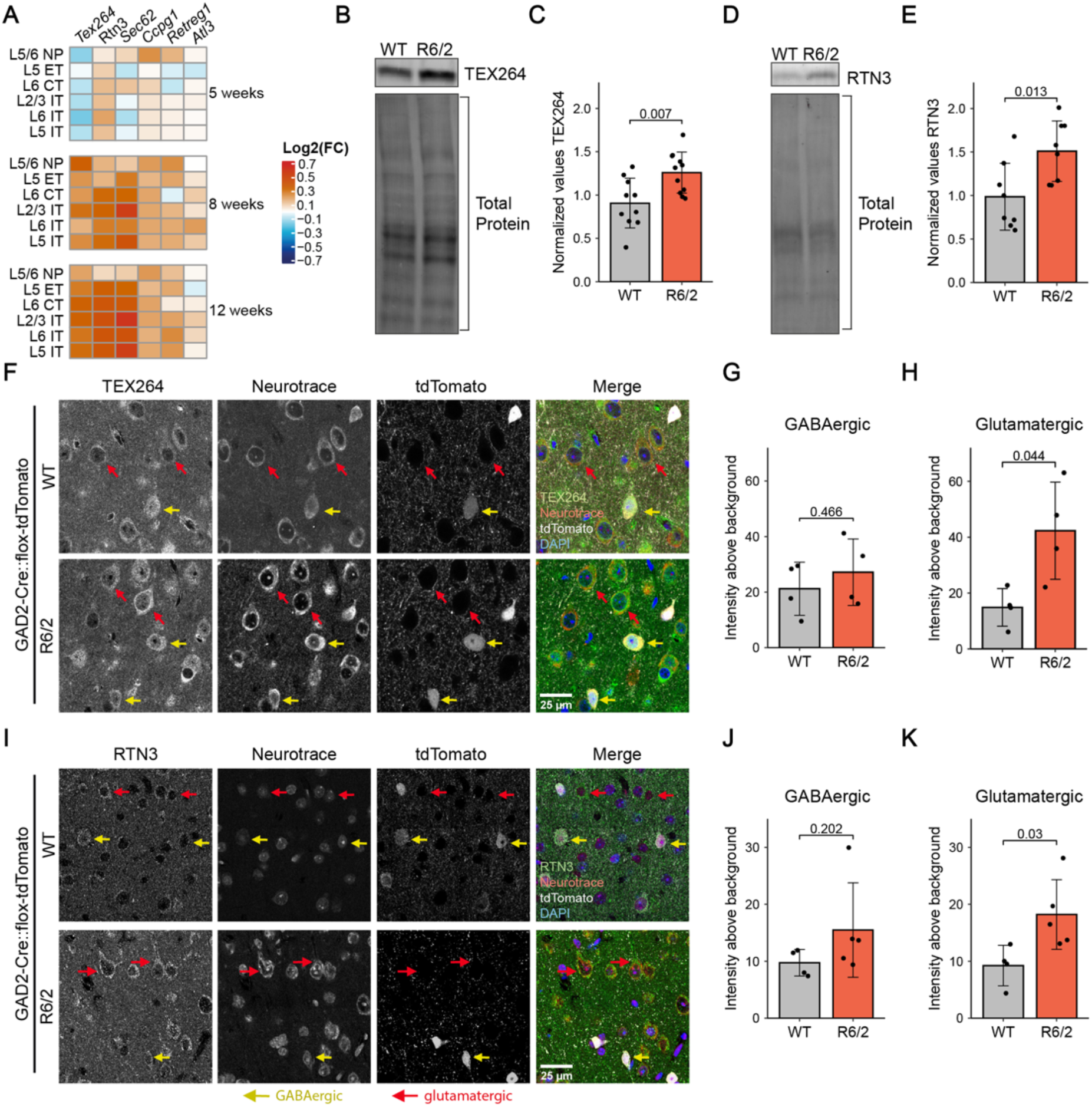
Increased expression of ER-phagy receptors in HD mouse cortex. **A**, Heat map of ER-phagy receptors expression in glutamatergic neuronal subclasses of R6/2 mice. **B**, Western blot for TEX264 in cortical lysates from 12-week-old R6/2 mice. Total protein staining was used as loading control. **C**, Quantification of TEX264 protein levels in R6/2 and WT mice, normalized to total protein. n=10 WT and 11 R6/2 mice. Unpaired two-sided t-test. **D**, Western blot for RTN3 in cortical lysates from 12-week-old R6/2 mice. Total protein staining was used as loading control. **E**, Quantification of RTN3 protein levels in R6/2 and WT mice, normalized to total protein. n=8 WT and 8 R6/2 mice. Unpaired two-sided t-test. **F**, Immunostaining for TEX264 on brain sections of 12-week-old R6/2 and control mice. NeuroTrace was used to detect neurons, GABAergic neurons were genetically labeled with tdTomato. Nuclei were stained with DAPI. Yellow arrows point to GABAergic neurons, red arrows to glutamatergic neurons. **G-H**, Quantification of TEX264 immunostaining in GABAergic (G) and glutamatergic (H) neurons. n=4 WT and 4 R6/2 mice. Unpaired two-sided t-test. **I**, Immunostaining for RTN3 on brain sections of 12-week-old R6/2 and control mice. NeuroTrace was used to detect neurons, GABAergic neurons were genetically labeled with tdTomato. Nuclei were stained with DAPI. Yellow arrows point to GABAergic neurons, red arrows to glutamatergic neurons. **J-K**, Quantification of RTN3 immunostaining in GABAergic (J) and glutamatergic (K) neurons. n=4 WT and 5 R6/2 mice. Unpaired two-sided t-test. P-values are indicated above the bars.

To test whether the upregulation of ER-phagy components also occurred at the protein level, we performed western blots for two of the ER-phagy receptors whose transcripts were strongly elevated, TEX264 and RTN3. Levels of both proteins were significantly higher in cortical lysates of R6/2 mice compared to WT controls (Fig. 6B-E). We next asked whether the upregulation was cell class-specific. To this end, we genetically labeled GABAergic neurons by crossing R6/2 mice to the GABAergic-specific GAD2-Cre line (Taniguchi et al., 2011) and the Cre-dependent tdTomato reporter (Madisen et al., 2010). Glutamatergic neurons were identified as Neurotrace-positive, tdTomato-negative cells. Immunostaining revealed significantly increased expression of TEX264 in glutamatergic neurons of R6/2 mice, while expression in GABAergic neurons was not significantly different from WT controls (Fig. 6F-H). RTN3 showed a similar pattern with significantly elevated expression in glutmatergic, but not GABAergic neurons of R6/2 mice (Fig. 6I-K). Altogether, these results demonstrate increased expression of ER-phagy receptors in HD-vulnerable neurons of R6/2 mice.

Upregulation of ER-phagy receptors suggested that mHTT might cause an increase in ER-phagy. We tested this in mHTT-expressing HEK293T cells and primary neurons with the help of a tandem ER-phagy reporter (Chino et al., 2019). The reporter consists of a signal sequence for targeting to the ER, two fluorescent proteins placed in tandem, RFP and GFP, followed by the KDEL ER-retention sequence (ssRFP-GFP-KDEL). At neutral pH in the ER, it emits fluorescence in both red and green channels. When ER-phagy is induced, the reporter is delivered to lysosomes and cleaved, resulting in quenching of GFP fluorescence in the acidic lysosomal environment, while the more stable RFP fragment remains fluorescent (Fig. 7A). We generated a stable HEK293T cell line with the ssRFP-GFP-KDEL reporter and transfected it with HTT-exon1 constructs containing a normal (HTTQ25-His) or pathologically elongated polyQ tract (HTTQ72-His). The numbers of red puncta per cell were quantified to assess accumulation of the RFP fragment in lysosomes as an indicator of ER-phagy induction (Supplementary Fig. S7A-B). Culturing the cells in Earle’s Balanced Salts Solution (EBSS) starving medium, known to induce autophagy, served as a positive control. Cells expressing the pathological HTTQ72-His protein had significantly higher numbers of red puncta per cell, indicative of increased ER-phagy, than control cells (Supplementary Fig. S7A-B).

**Figure 7.**
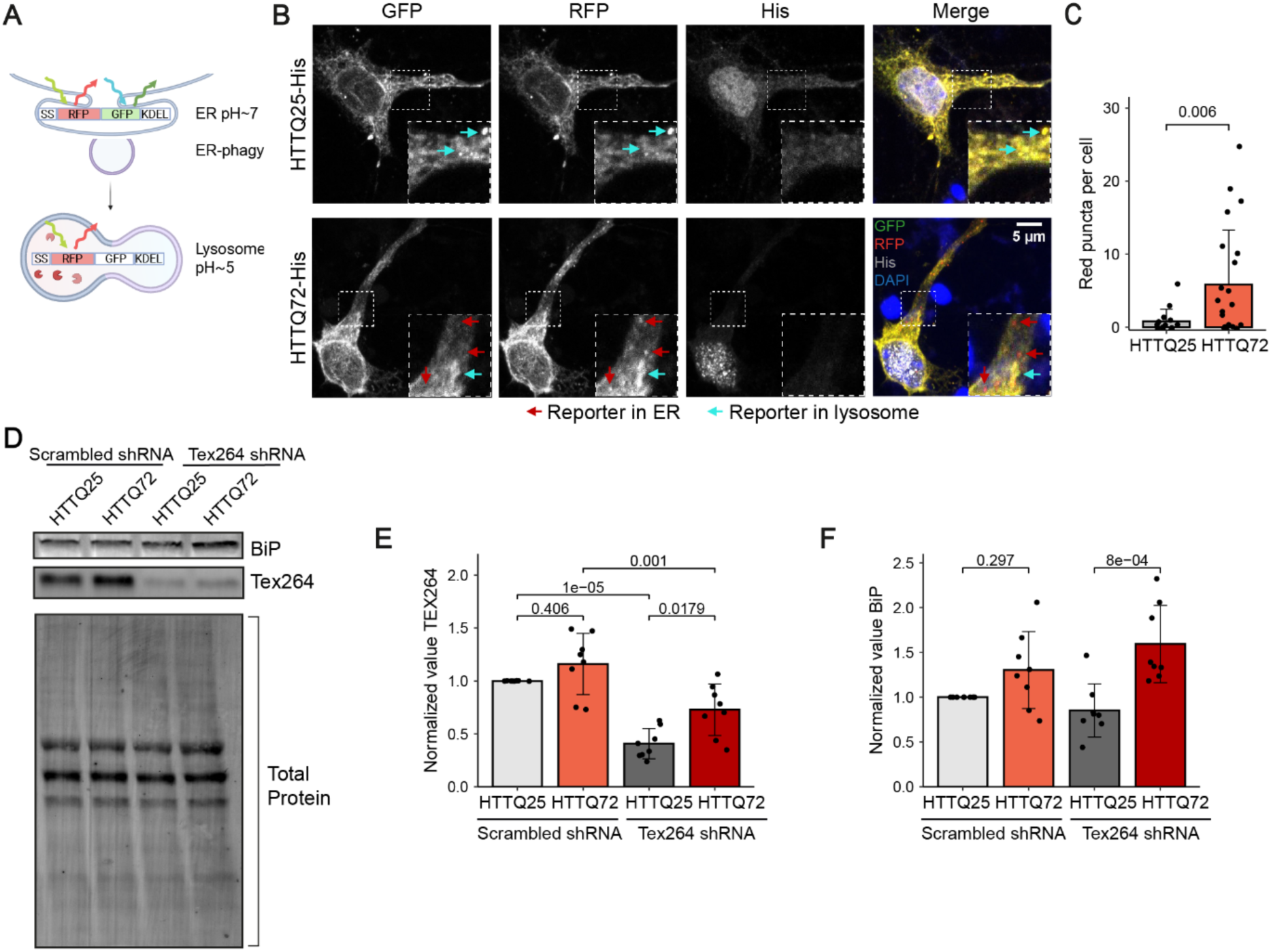
ER-phagy is increased in the presence of mHTT. **A**, Principle of the ER-phagy sensor. **B**, Representative images of primary neurons co-transfected with the ER-phagy sensor and the indicated HTT constructs. HTT was detected by staining against His-tag. Insets show higher magnifications of the areas marked by dashed boxes. Cyan and red arrows point to ssRFP-GFP-KDEL reporter in the ER and in lysosomes, respectively. **C**, Quantificaton of red puncta corresponding to reporter inside lysosomes in the indicated conditions. n=15 HTTQ25-His neurons and 22 HTTQ72-His neurons from 7 independent cultures. Unpaired two-sided t-test. **D**, Western blot for BiP and TEX264 in primary neuron lysates. Total protein staining was used as loading control. **E**, Quantification of TEX264 knockdown in the indicated conditions. **F**, Quantification of BiP protein levels in the indicated conditions. n=8 independent experiments. Two-way ANOVA, followed by Tukey’s post-hoc test. P-values are indicated above the bars.

We next asked whether mHTT also induces ER-phagy in neurons. Primary murine cortical neurons were co-transfected with the ER-phagy reporter ssRFP-GFP-KDEL and non-pathological HTTQ25-His or pathological HTTQ72-His constructs, which forms primarily nuclear aggregates in neurons (Voelkl et al., 2023) (Fig. 7B). The number of red puncta per cell was significantly higher in pathological HTTQ72-His compared to control HTTQ25-His cells, indicating that nuclear mHTT aggregates are associated with increased ER-phagy in neurons (Fig. 7C).

mHTT is known to promote ER stress and activate the UPR (Leitman et al., 2013; Vidal et al., 2012), which is supported by our findings of upregulated ER homeostasis genes in HD mice (Supplementary Fig. 5D). ER-phagy is a crucial pathway to relieve ER stress (Chino and Mizushima, 2020; Yang et al., 2021). ER-phagy can be induced through the UPR and in turn acts to prevent excessive activation of the UPR (Hanquier et al., 2023; Smith et al., 2018). We therefore hypothesized that in HD, ER-phagy might represent a compensatory mechanism aimed at reducing mHTT-dependent ER stress. To explore this possibility, we quantified the amount of BiP, a well-established UPR target gene (Hendershot, 2004). We reasoned that in mHTT-expressing neurons with increased ER-phagy the levels of BiP should remain low, whereas under conditions of impaired ER-phagy the levels of BiP should increase. Primary murine cortical neurons were co-transduced with lentiviral vectors encoding either normal HTTQ25-myc or pathological HTTQ72-myc, and either a scrambled shRNA or a shRNA targeting the ER-phagy receptor *Tex264* transcripts to impair ER-phagy (Fig. 7D-E). In neurons with scrambled shRNA, expression of pathological HTTQ72-myc only caused a slight elevation of BiP levels, which was not statistically significant (Fig. 7D, F). In contrast, expression of HTTQ72-myc in *Tex264* knockdown neurons resulted in a significant increase in BiP, indicating aggravated mHTT-dependent ER-stress in conditions of impaired ER-phagy (Fig. 7D, F). In summary, our findings suggest that ER-phagy is increased in response to mHTT expression, and alleviates mHTT-induced ER stress.

## Discussion

### Cell type-resolved transcriptomic disease signatures in HD mouse cortex

Here, we explored transcriptional disease signatures in the motor cortex of HD mice. Investigation of the cortical tissue enabled us to compare vulnerable (glutamatergic) and resistant (GABAergic) neuronal cell types within the same sample. We show that glutamatergic neurons exhibit more prominent transcriptomic changes, consistent with a large body of neuropathological data pointing to pyramidal cell degeneration in patients (Waldvogel et al., 2012). Among the glutamatergic cell types, IT cells of different cortical layers emerged as the populations with most significant transcriptomic dysregulation. This is in agreement with the crucial role of corticostriatal miscommunication and deafferentation in HD, as IT cells constitute the main source of corticostriatal projections (McColgan et al., 2020).

In contrast to previous single-cell transcriptomic studies in HD models, we employed a temporally-resolved approach and investigated not only advanced, but also early stages of pathology, allowing us to probe transcriptomic changes in presymptomatic animals. Surprisingly, we observed lower than WT pseudotime values of R6/2 nuclei at 5 weeks, while at 8 and 12 weeks the values were higher than WT. A possible explanation is that expression of mHTT might trigger compensatory responses that efficiently counteract the effects of mHTT at the early presymptomatic stage of 5 weeks, but later fail, potentially contributing to the onset of symptoms. For example, the transitory downregulation of ribosomal genes at the early stage might represent an attempt to reduce protein synthesis in order to relieve mHTT-induced proteotoxis stress. The pronounced transcriptional changes that take place in the R6/2 brain between 5 and 8 weeks are consistent with the marked proteome remodeling between these time points, which we observed previously (Hosp et al., 2017). Furthermore, our findings are in line with a recent spatial transcriptomics study, which revealed a biphasic regulation of the OXPHOS pathway in the R6/2 mouse cortex, with downregulation at early and upregulation at later disease stages (Burns et al., 2025). Consistently, oxidative phosphorylation genes were among the prominent biphasically regulated categories also in our dataset.

Our KEGG pathway analysis revealed downregulation of many genes involved in neuron-specific processes, such as axon guidance and synaptic function. This is in agreement with transcriptomic and proteomic analyses in HD mice and HD patient-derived induced pluripotent stem cells that point to changes in neuronal development and synaptic function (HD iPSC Consortium, 2017). Moreover, a large body of experimental data supports both neurodevelopmental defects and synaptic impairments in HD (Barnat et al., 2020; Braz et al., 2022; Molina-Calavita et al., 2014; Veldman and Yang, 2017).

Another interesting observation in our study was the general decline in the expression of cell subclass-specific genes. Loss of endogenous transcriptomic cellular identities has been recently observed also in striatal cell populations of HD mice and patients, as well as in oligodendroglia across several brain regions (Lim et al., 2022; Malaiya et al., 2021; Matsushima et al., 2023). Accordingly, our histological analyses revealed that cortical Sst and VIP interneurons display a partial loss of immunoreactivity for their specific marker proteins in HD mice (Voelkl et al., 2022). The study by Malaiya et al. attributed the compromised cell identity to the altered interaction of HTT with the polycomb repressor complex 2 (PRC2). This interaction is disturbed by the HTT mutation, liberating PRC2 from the complex with HTT and facilitating its binding to the promoters of neuronal genes, which results in increased repression of these genes (Malaiya et al., 2021). Considering our observation of reduced expression levels of multiple neuronal transcription factors in glutamatergic neurons, it is intriguing to speculate that PRC2 might play a role in the impaired maintenance of neuronal cell identity across brain regions, and might represent an attractive target for potential interventions. However, the precise details of this mechanism remain to be deciphered.

Interestingly, we find largely similar categories of DEGs in glutamatergic as well as in GABAergic neurons, with the changes being less significant in GABAergic cells. This is compatible with the scenario where all neuronal subclasses undergo qualitatively similar molecular changes, which are however delayed and/or less pronounced in the GABAergic cell types. The underlying causes of these differences is an interesting question for future investigations. Recent studies suggest that differential vulnerability cannot be fully explained by the varying expression levels of the disease-causing pathological proteins, but is likely due to intrinsic molecular differences between the cell types (Praschberger et al., 2023). One factor contributing to the resilience of interneurons to degeneration might be the specific repertoire of protein quality control factors expressed by these cells (Fu et al., 2019). Indeed, our analyses revealed that GABAergic neurons show higher baseline expression values of the neuroprotective turquoise module, which contains a number of proteostasis-related hub genes (Fig. 4D-E and 5).

While we did not focus on glial cells in this study, our results complement several reports investigating glial pathology in HD. Thus, we observed increased cell numbers (Fig. 1B-C), transcriptomic shifts (Fig. 1D-E) and reduced marker gene expression (Fig. 3E) in the oligodendrocyte and astrocyte populations. These findings are in line with previous histological analyses in human HD patients (Myers et al., 1991) and with recent transcriptomic studies in mice implicating oligodendrocytes and astrocytes in HD pathogenesis (Al-Dalahmah et al., 2020; Lim et al., 2022).

### Role of ER-phagy in HD pathogenesis

ER homeostasis-related genes were highly upregulated in the pathway analysis and gene network analysis, and occupied central positions in the gene network of R6/2 mice. Notably, *Hspa5* and *Xbp1*, which are a key activator and regulator of the UPR, respectively, were among the earliest upregulated genes and remained elevated at later time points, indicating activation of the UPR and chronic ER stress (Supplementary Fig. S5D). The occurrence of ER stress in the context of HD is well established and might be caused at least in part by the sequestration of valosin-containing protein (VCP/p97), a chaperone important for the process of ER-associated degradation, by mHTT oligomers. This impairs protein folding in the ER and causes accumulation of unfolded and misfolded proteins in the ER lumen (Leitman et al., 2013). Prolonged ER stress leading to chronic UPR activation can be deleterious and drive cell death (Lindholm et al., 2006). Neurons failing to restore ER homeostasis may therefore be particularly prone to neurodegeneration.

The consistent signatures of ER stress prompted us to investigate the role of ER-phagy as a previously unexplored component of stress response in HD. So far, the involvement of ER-phagy in neurodegenerative diseases has been understudied and direct evidence of its involvement is sparse (Hill et al., 2023). A recent study showed that ER-phagy mediated by RETREG1 (FAM134B) is capable of clearing overexpressed α-synulcein from the ER, exerting a neuroprotective effect (Kim et al., 2023). Here we found that several ER-phagy receptors, including TEX264 and RTN3, were upregulate on the transcript and protein level in the cortex of HD mice. Interestingly, the upregulation occurred selectively in glutamategic neurons, which are more vulnerable to HD. Furthermore, mechanistic experiments in cellular models demonstrated increased ER-phagy in the present of mHTT. ER-phagy might ameliorate ER stress and restore ER homeostasis by removing the proteotoxic burden. This hypothesis is supported by our finding that knockdown of *Tex264* in the presence of mHTT exacerbated ER stress, as indicated by heightened BiP expression. Notably, TEX264 previously was shown to alleviate ER stress in the context of neuroinflammation (Wang et al., 2022), raising the possibility of a similar role of TEX264 in HD. In summary, our data suggest that ER-phagy might serve as a protective mechanism in HD aimed at restorating ER homestasis. Further studies should investigate whether enhancing ER-phagy in HD could be beneficial and delay neurodegeneration.

## Materials and Methods

### Animals

Animal experiments were approved by the Government of Upper Bavaria, Germany (animal protocols ROB-55.2-2532.Vet_02-20-05 and ROB-55.2-2532.Vet_02-23-213) and performed in accordance with the relevant guidelines and regulations. Mice were housed in SPF-free environment with ad libitum access to food and water. Transgenic R6/2 line (Mangiarini et al., 1996); JAX stock #002810) was maintained by breeding hemizygous R6/2 males with the female F1 progeny of a cross between CBA (Janvier Labs) and C57BL/6 (Janvier Labs) mice. Length of the CAG repeat tract in the R6/2 mice used for experiments was determined by Laragen, Inc., and averaged 204 ± 5.85 (SEQ CAG No., mean ± sd). tdTomato (Madisen et al., 2010), JAX stock #007909) and GAD2-Cre mice (Taniguchi et al., 2011), JAX stock #010802) were kept on C57BL/6 background. For genetic tracing experiments, hemizygous R6/2 males were crossed to mice heterozygous for the Cre allele and homozygous for the tdTomato reporter allele. Mice of either sex were used for all experiments.

### Plasmids

For expression via transfection, pcDNA3.1 HTT-exon1-Q25-His, HTT-exon1-Q72-His (cloned from pPGK HTT-exon 1-Q25, HTT-exon 1-Q72) (Jeong et al., 2011) and pCW57-CMV ssRFP-GFP-KDEL (Chino et al., 2019) were used.

For lentivirus packaging, psPAX2 and pcDNA3.1-VSV-G (kindly provided by Dieter Edbauer) were used. For lentiviral expression, HTT fragments from pPGK HTT-exon1-Q25-myc, pPGK HTT-exon1-Q72-myc (Jeong et al., 2011) were cloned into the pLenti vector (Zhang et al., 2011). For TEX264 knockdown, pRP[shRNA]-EGFP-U6>Scramble_shRNA and pLV[shRNA]-EGFP-U6>mTex264[shRNA#1] (constructed by VectorBuilder) were used.

#### Tissue dissection

Tissue was harvested from 12 R6/2 and 14 WT mice in total: 4 R6/2 and 4 WT at 5 weeks; 4 R6/2 and 6 WT at 8 weeks; and 4 R6/2 and 4 WT at 12 weeks. Tissue isolation and preparation of single nuclei were conducted with WT and R6/2 littermates in parallel to reduce the possibility of batch effects. To reduce the variance originating from single animals, for each replicate motor cortices were sampled from two animals of the same genotype.

Buffers for cortex dissection were adapted from (Saunders et al., 2018). Mice were anesthetized by i.p. injections of 200 mg/kg Ketamine (MEDISTAR) and 40 mg/kg Xylazine (MEDISTAR). Mice were perfused by injection of ice-cold cutting buffer (110 mM NaCl, 2.5 mM KCl, 10 mM HEPES [Biomol, 05288-100], 7.5 mM MgCl_2_, and 25 mM glucose, 75 mM sucrose, pH=7.4). All following procedures were performed at 4°C. The brain was extracted and cut at a thickness of 300 µM in cutting buffer on a Leica VT 1200S vibratome at 0.2 mm/s. Brain slices were transferred into dissection buffer (82 Na_2_SO_4_, 30 K_2_SO_4_, 10 HEPES, 10 glucose and 5 MgCl_2_), and the primary motor cortex was microdissected.

#### Nuclei isolation and snRNA-seq

Tissue samples from 2 mice were pooled together for nuclei isolation. The protocol for nuclei isolation was adapted from (Mathys et al., 2019) and (Wertz et al., 2020). Motor cortices were transferred to a 2 mL KIMBLE tissue douncer with 600 µL homogenization buffer (320 mM sucrose, 5 mM CaCl_2_, 3 mM Mg(CH_3_COO)_2_, 10 mM Tris HCl [pH 7.8] [Thermo Fisher Scientific, T2569], 0.1 mM EDTA [pH 8.0] [Thermo Fisher Scientific, 15575020], 0.1% NP-40 [Sigma-Aldrich, NP40S], 1 mM β-mercaptoethanol [Sigma-Aldrich, M-7522], and 0.4 U/mL SUPERaseIn RNase Inhibitor [Invitrogen, AM2694]). The tissue was dounced 15 times with a loose and 15 times with a tight pestle. The homogenized tissue was passed through a 20 µm cell strainer. The filtered solution was mixed with an equal volume of working solution (50% OptiPrep density gradient medium [Sigma-Aldrich, D1556], 5 mM CaCl_2_, 3 mM Mg(CH_3_COO)_2_, 10 mM Tris HCl [pH 7.8], 0.1 mM EDTA [pH 8.0], and 1mM β-mercaptoethanol). The mixed solution was transferred to a Sorenson microcentrifuge tube and underlayed first with a 30% Optiprep solution (134 mM sucrose, 5 mM CaCl_2_, 3 mM Mg(CH_3_COO)_2_, 10 mM Tris HCl [pH 7.8], 0.1 mM EDTA [pH 8.0], 1 mM β-mercaptoethanol, 0.04% NP-40, and 0.17 U/mL SUPERase In RNase Inhibitor) and then with a 35% Optiprep solution (96 mM sucrose, 5 mM CaCl_2_, 3 mM Mg(CH_3_COO)_2_, 10 mM Tris HCl [pH 7.8], 0.1 mM EDTA [pH 8.0], 1 mM 2-mercaptoethanol, 0.04% NP-40, and 0.12 U/mL SUPERase In RNase Inhibitor). The gradient was centrifuged for 20 min at 3000 x g on a SH 3000 swing rotor. 200 µL of interface between the 35% and 30% Optiprep solution containing the nuclei was collected. For washing, the nuclei solution was diluted in 1000 µL 2% Albumin Fraction V (BSA) [Karl Roth, 8076.4] in PBS, thoroughly mixed and centrifuged for 4 min at 300 x g. The supernatant was removed and the washing step repeated once. The nuclei were counted and the nuclei solution diluted to 300 nuclei/µL in 2% BSA in PBS. A volume targeting for 12,000 nuclei was loaded onto a Chip G (10X Genomics, PN-1000120) on a 10X Chromium microfluidics controller (10X Genomics). Sequencing libraries were prepared with the Chromium Single Cell 3’ reagent kit v3.1 (10X Genomics, PN-1000121) according to manufacturer’s protocol. Library indices were added with the Single Index Kit T (10X Genomics, PN-1000213).

Sequencing was performed on a NovaSeq6000 at Core Facility Genomics of the Helmholtz Zentrum München. BCL files were demultiplexed to FASTQ files with Cell Ranger v4.0.0 (10X genomics) using mkfastq(). Count matrices were generated by aligning FASTQ files to the mouse GRCm38 reference genome (NCBI: GCF_000001635.20) with cellranger count() with intronic reads being included.

#### Data Analysis: Quality control and cell type annotation

Seurat v4.0.1 was used for subsequent data cleaning, normalization, integration into reference genome, visualization and calculation of DEGs. Cells with a higher UMI count than 30,000, with a lower UMI count than 250 or expressing less than 200 genes were removed. Cells with more than 2% mitochondrial genes were removed. Following annotation of nuclei, non-neuronal nuclei with less than 500 UMI and neurons with less than 1,200 UMI were removed. Normalization of UMI was done with NormalizeData() with a scale factor of 10,000 and “LogNormalize” as normalization method. The top 4,000 highly variable genes were detected with FindVariableGenes(). For PCA, data was scaled with ScaleData() and PCA was done using RunPCA() with npcs = 35. UMAP was done with RunUMAP() with dims=30.

For annotation, our data set was integrated into the mouse primary motor cortex reference atlas published by (Yao et al., 2021). We used “snRNAseq 10x v3 data set B” as a reference, as it was generated with an improved nuclei isolation protocol and detected the most median genes per nucleus. We integrated our data set into the reference atlas using the FindTransferAnchors() function with canonical correlation analysis as reduction method before finding common anchors. Cluster annotation for each nucleus was predicted using TransferData(). Cell subclass annotation was retrieved by summarising cell types according to the BICCN annotation (Yao et al., 2021). Doublets were removed using doublet finder package v2.0.3 (McGinnis et al., 2019) assuming 6.1% of doublets.

#### Differential abundance testing

Differential abundance testing was performed using the *miloR* package v1.4.0.0 (Dann et al., 2022). A k-nearest neighbor graph was constructed with buildGraph() with k = 50 and d = 20. Neighborhoods were defined using makeNhoods(), with randomly sampled neighborhoods set to prop = 0.1. The contrast was set as "R6/2 vs WT". Significantly different neighborhoods between conditions were identified using testNhoods() and defined as having an FDR-corrected p-value < 0.1. The plotDAbeeswarm() function was used to generate swarm plots, and plotNhoodGraphDA() was used to plot neighborhood graphs.

#### Trajectory Analysis

Trajectory analysis was performed by subplotting cell subclasses on a UMAP. Larger subclasses (L5 IT and L6 IT) were subplotted by cell type. Trajectories and pseudotime values were inferred using the slingshot() function from the *Slingshot* package v2.4.0 (Street et al., 2018), applied to the UMAP coordinates. WT nuclei were set as the starting point and R6/2 nuclei as the endpoint with curve extension being allowed, no breaks, and a smoothing approximation across 150 nuclei.

#### Differentially expressed genes and gene enrichment analysis

Cell subclass-specific DEGs were identified using FindMarkers() from Seurat v4.0.1 with the Wilcoxon rank-sum test. DEGs were defined as having an adjusted p-value < 0.05, |log2FC| > 0.2 and being expressed in at least 5% of nuclei of the respective cell subclass. To validate the results of the Wilcoxon rank-sum test at 8 weeks and 12 weeks, pseudobulk counts were generated for each cell subclass and each sample. Afterwards, bulk DEG analysis was performed with DESeq2 (Love et al., 2014) between WT and R6/2 with combined pseudobulk counts from 8 weeks and 12 weeks.

KEGG gene enrichment analysis was performed with the clusterProfiler package v4.10.0 (Wu et al., 2021). DEGs of the specific cell subclass were taken into consideration and enrichment analysis was performed separately for downregulated and upregulated genes. Enriched terms were defined as having a Benjamini-Hochberg-corrected p-value <0.05.

#### scWGCNA

WGCNA was performed using the scWGCNA package v0.1.0 (Feregrino and Tschopp, 2022). scWGCNA was conducted using all neuronal clusters. For pseudonuclei generation, nuclei were subset by genotype and time point. Pseudonuclei were generated using 20% of nuclei as seeds and 15 nearest neighbors in UMAP space (dims=1:2). scWGCNA was performed considering the top 2,000 highly variable genes and DEGs that occurred in at least one time point in at least 2 cell subclasses. Genes expressed in less than 1% of cells were filtered out. WGCNA threshold was chosen using pickSoftThreshold() with a cut-off power of 0.9 for a signed network. WGCNA was performed iteratively until all genes were assigned to a module and small modules as described in (Feregrino and Tschopp, 2022). Module eigenegene expression for nuclei was calculated with moduleEigengenes() of the WGCNA package (Langfelder and Horvath, 2008). Gene network representations were plotted using the igraph package v1.4.1 with Fruchterman-Reingold representation. The top 10% of hub genes from modules with a module-trait relationship > |0.15| for at least one time point were plotted. The vertex sizes are proportional to module eigengene values and edges represent the top 20% highest absolute adjacencies.

P-values for the over-representation of neuronal essential genes and mHTT modifier genes in WGCNA gene modules were calculated using one-sided Fisher’s exact test, following the same principle of KEGG pathway enrichment analysis.

#### HEK cell culture

HEK-293T cells were cultured in HEK medium (88% Dulbecco’s Modified Eagle Medium [DMEM], with high glucose [Thermo Fisher Scientific, 41965062], 10% fetal bovine serum [FBS] [Sigma-Aldrich, F7524], 1% non-essential amino acids [Gibco, 11140050], 1% Penicillin-Streptomycin [Thermo Fisher Scientific, 10378016]). Stable ssRFP-GFP-KDEL HEK cells were maintained in the same medium supplemented with 0.5 µg/mL Puromycin (MP Biomedicals, 11497610). Cells were cultured at 37 °C in a humidified incubator with 5% CO₂ and passaged before reaching 90% confluence. HEK cells were split every 2–3 days. For splitting, cells were washed once with PBS, incubated with trypsin (Sigma-Aldrich, T4299) for 5 min at 37 °C, resuspended in 5 mL HEK medium, and centrifuged for 3 min at 300 × g. The supernatant was discarded and the pellet resuspended in 10 mL HEK medium. 1 mL of this suspension was transferred to a new flask for 1:10 splitting. HEK cells were used up to passage 25 and stable ssRFP-GFP-KDEL HEK cells were used up to passage 20.

#### Transfection of HEK cells

HEK cells were seeded at 30% confluence in 24-well plates using antibiotic-free HEK medium. Transfection was performed when the density reached 70–80% confluence using 0.5 µg of DNA diluted in 100 µL Opti-MEM (Gibco, 31985070), mixed with 1.5 µL Lipofectamine LTX (Thermo Fisher Scientific, 15338030) per well. After 30 min incubation at room temperature, the DNA–Lipofectamine mixture was added dropwise. The medium was replaced after 48 h. For selection of stably transfected cells expressing the ssRFP-GFP-KDEL construct, 1 µg/mL Puromycin was added and refreshed every 48 h. After 10 days of selection, cells were dissociated and diluted stepwise to generate suspensions ranging from 100 to 3,125 cells/mL. From each dilution, 5 × 100 µL aliquots were plated into 96-well plates. Single-cell colonies were visible after ∼10 days and were transferred to 6-well plates for expansion. From this point on, stable ssRFP-GFP-KDEL HEK cells were cultured in HEK medium with 0.5 µg/mL Puromycin. For expression of ssRFP-GFP-KDEL, 0.5 µg/mL of doxycycline (Sigma-Aldrich, D9891) was added to the medium 24 h before fixation of the cells with 4% PFA (Thermo Fisher Scientific, J19943.K2) for 20 min.

Transfection for lentivirus production was done in HEK cells in T175 flasks at 80% confluence. A plasmid mix (18.6 μg transfer plasmid, 11 μg psPAX2, 6.4 μg pMD2.G) was mixed with 108 μl TransIT-Lenti reagent (Mirus Bio, MIR 6604) in 1,398 μl DMEM, and incubated 20 min at room temperature. The solution was added dropwise to the HEK cells. 24 h after transfection, the medium was removed and 30 mL fresh HEK medium added. 48 h after medium change, viral supernatant was collected and centrifuged at 1,200 x g for 20 min, filtered (0.45 μm), and concentrated with Lenti-X Concentrator (Takara Bio, 631231) overnight at 4 °C according to the manufacturer’s protocol. The solution was centrifuged at 1,500 x g for 45 min at 4 °C. The supernatant was removed and the pellet dried for 5 min at room temperature. Viral pellets were resuspended in PBS, incubated for 4 h at 4 °C, and stored at −80 °C.

#### Primary cortical murine cultures

Pregnant CD-1 females with embryos at stage E15.5 were sacrificed through cervical dislocation. Embryos were dissected, transferred into ice-cold PBS, decapitated, and transferred into dissection medium (Hanks’ Balanced Salt Solution [HBSS] [Thermo Fisher Scientific, 24020-117], 1% Pen/Strep, 1% HEPES, 1% MgSO₄). Brains were taken out, meninges removed, and neocortices dissected. The neocortices were incubated in pre-warmed trypsin with 0.05% DNase I (Thermo Fisher Scientific, EN0521) for 15 min at 37 °C, and trypsin was afterwards inactivated with Neurobasal medium containing 5% FBS. After washing, tissue was resuspended in neuronal culture medium (96% Neurobasal medium [Thermo Fisher Scientific, 21103049], 2% B27-supplement [Thermo Fisher Scientific, 17504044], 1% Pen/Strep, 1% L-Glutamine [Thermo Fisher Scientific, 25030123]). The neocortices were dissociated by pipetting them 15 times up and down, and cells were pelleted by centrifugation at 130 x g for 5 min. The cells were resuspended in neuronal culture medium at a concentration of 250,000 cells/mL. Cells were plated at 125,000/well in 24-well plates on glass coverslips or 500,000/well in 6-well plates, both coated with poly-L-lysine (Sigma-Aldrich, P7886) and laminin (Gibco, 23017015), and maintained at 37 °C with 5% CO₂.

#### Transfection of primary cortical murine neurons

Primary cortical neurons were transfected on day *in vitro* (DIV) 7 using NeuroMag Transfection Reagent according to the manufacturer’s protocol. In short, for each well to be transfected, 1 µg of ssRFP-GFP-KDEL plasmid and 2 µg of either HTTQ25-his or HTTQ72-his plasmid were added to 100 µL of Neurobasal medium and mixed with 2 µg of NeuroMag (OZ Biosciences, NM50200). The solution was mixed, incubated for 15 min at RT, added dropwise to wells and incubated on a magnetic plate for 15 min at 37 °C with 5% CO₂. For the expression of ssRFP-GFP-KDEL, 0.5 µg/mL doxycycline was added to the medium 24 h before fixation of cells with 4% PFA for 20 min.

#### Transduction

Primary cortical neurons were transduced on DIV 5 with lentivirus encoding HTTQ25-myc or HTTQ72-myc. On DIV 7, lentivirus encoding either an shRNA targeting *Tex264* or a scrambled control was added. For each well of a 6-well plate, 200 µL of pre-warmed neuronal culture medium was mixed with 0.5–1 µL of viral suspension, depending on viral titer. The entire 200 µL mixture was added directly to the wells.

#### Immunocytochemistry

Fixed cells were washed with PBS for 5 min, then incubated with 50 mM NH₄Cl in PBS for 10 min. Permeabilization was performed with 0.25% Triton X-100 (Karl Roth, 6683.1) in PBS for 5 min, followed by one PBS wash. Cells were placed in a humid chamber and blocked for 30 min in blocking solution (2% BSA, 4% normal donkey serum [abcam, ab7475] in PBS). Primary mouse anti-His antibody (1:500, Diazol, Dia-900-100) was diluted in blocking solution, and cells incubated with 50 µL of primary antibody solution for 1 h. Cells were then washed 3× with PBS. Secondary antibody donkey anti-mouse Alexa Fluor 647 (1:250, Jackson, 715-065-140) was diluted in blocking solution with and cells were incubated with 50 µL of secondary antibody solution for 30 min. Cells were washed 3× 5 min with PBS. During the second wash, DAPI (1:1000, Sigma-Aldrich, D9542) was added to the solution. Coverslips were rinsed with distilled H₂O and were mounted using 10 µL ProLong Glass Antifade Mountant (P36984; Invitrogen) and stored at 4 °C until imaging. Images were acquired from randomly selected cells using a Leica TCS SP8 confocal microscope.

#### Immunostainings of brain sections

Mice were transcardially perfused with PBS for 5 min, then with 4% PFA in PBS for 5 min at 3-3.5 mL/min under ketamine/xylazine anesthesia. Brains were dissected and incubated in 4% PFA overnight at 4 °C. Brains were cut into 70 μm thin sections on a vibratome. All following steps were carried out with protection from light to prevent bleaching of fluorophores and with gently shaking. Sections were washed with PBS, incubated with 0.05% Triton X-100 in PBS for 15 min and antigen retrieval was performed by incubating the sections in 10 mM trisodium citrate pH 6 with 0.05% Tween 20 (Bio-Rad, 1706531) for 15 min at 80 °C at 300 rpm on an Eppendorf ThermoMixer. After letting the sections cool down, they were incubated in blocking solution (0.2% BSA, 5% normal donkey serum [abcam, ab7475], 0.2% lysine [Thermo Fisher Scientific, J62225.36] and 0.2% glycine [Thermo Fisher Scientific, A13816.36] in PBS) for 1 h. Sections were incubated with primary antibodies in blocking solution overnight at 4 °C. The following antibodies were used: rabbit anti-TEX264 (1:100, 25858-1-AP, Proteintech) and rabbit anti-RTN3 (1:200, BS-7644R, Thermo Fisher Scientific). Afterwards, sections were washed 3 times for 10 min with PBS at RT, followed by incubation with secondary antibody solution for 1 h at RT with donkey anti-rabbit Alexa Fluor 488 (1:250, 711-545-152, Jackson) in blocking solution with 1:500 NeuroTrace 640/660 (Invitrogen, 10266952). Afterwards, sections were washed 3 times for 10 min with PBS and during the second wash, 1:1000 DAPI was added to the solution. Sections were mounted using ProLong Glass Antifade Mountant.

#### Western Blot

For preparation of mouse brain tissue lysates, mice were euthanized by cervical dislocation, and brains were extracted. The motor cortex was dissected on ice, flash-frozen in liquid nitrogen, and stored at −80 °C until use. For protein isolation, 100 μl of ice-cold lysis buffer (50 mM Tris–HCl, pH 7.5; 150 mM NaCl; 1% Triton X-100; 2 mM EDTA; protease inhibitor cocktail [Roche, 04693159001]; phosphatase inhibitor [Roche, 04906837001]) was added to each cortex tissue sample, which was then homogenized on ice using a tissue homogenizer until a uniform solution was obtained. Samples were subsequently incubated on ice for 30 min and stored at −20 °C until use.

For preparation of primary murine neuronal culture lysates, cells were washed with PBS, and 150 μl of ice-cold lysis buffer was added to each well, followed by a 10 min incubation at 4 °C. Cells were then detached using a cell scraper, and the lysate was collected. Samples were stored at −20 °C until use.

Protein samples were prepared, aiming for 20 μg of protein per sample, and 4× Laemmli sample buffer (Bio-Rad, 1610737) containing 10% β-mercaptoethanol was added. Samples were incubated at 95 °C, vortexed, and spun down afterwards. Proteins were separated by SDS-PAGE using 10% Criterion TGX Stain-Free gels (Bio-Rad, 5678035). Gels were activated and transferred to a 0.45 μm PVDF LF membrane (Thermo Fisher Scientific, 88518) using the Trans-Blot Turbo Transfer System (Bio-Rad) for 9 min at 2.5 A and 25 V. Images of total protein were taken with the ChemiDoc MP Imaging System (Bio-Rad). Membranes were blocked with 3% BSA in Tris-buffered saline with Tween20 (TBS-T) and incubated overnight at 4 °C with primary antibodies: BiP/GRP78 (mouse, 1:1,000, Proteintech, 66574-1-Ig) and TEX264 (rabbit, 1:1,000, Proteintech, 25858-1-AP). After washing, membranes were incubated for 1 h at room temperature with fluorescently labeled secondary antibodies: StarBright Blue 700 anti-mouse IgG (1:2,500, Bio-Rad, 12004159) and Alexa Fluor 647 anti-rabbit IgG (1:2,500, Invitrogen, A-21245). The signal of secondary antibodies was detected with the ChemiDoc MP Imaging System (Bio-Rad).

#### Statistics

Differences were considered statistically significant at p-value < 0.05, and for multiple comparisons, at an adjusted p-value < 0.05. In bar graphs, columns with error bars represent mean ± standard deviation. In box plots, boxes indicate median and interquartile range from the first (Q1) to the third quartile (Q3), and whiskers indicate Q1/Q3 -/+ 1.5 * interquartile range.

## Acknowledgements

We thank Ozgun Gökce and Christian Mayer for insightful discussions; Miguel da Silva Padilha for help with biochemistry; Marco Niestroj for help with immunostainings; Wing Han Liu, Magdalena Böhm and Sofia Delgado for mouse genotyping; and Marius Lemberg for critically reading the manuscript. This work was funded by the Max Planck Society for the advancement of Science, the Chan-Zuckerberg Initiative (to I.D.) and the Deutsche Forschungsgemeinschaft (DFG, German Research Foundation - SFB1451 project number 431549029-A09; SPP2453 project number 541742535; FOR5762 project number 531902955 and CECAD, EXC 2030-390661388 to I.D.). D.F. is recipient of an Add-on Fellowship for Interdisciplinary Life Science of the Joachim Herz Stiftung.

## Author contributions

I.D. conceived the project. D.F. and K.V. performed nuclei isolation and snRNA-seq experiments. D.F. performed immunostainings and cell biology experiments, analyzed the data and designed the figures. I.D. and R.K. supervised the project. D.F. and I.D. wrote the paper with input from all the authors.

## Supplementary figures

**Supplementary Figure S1.**
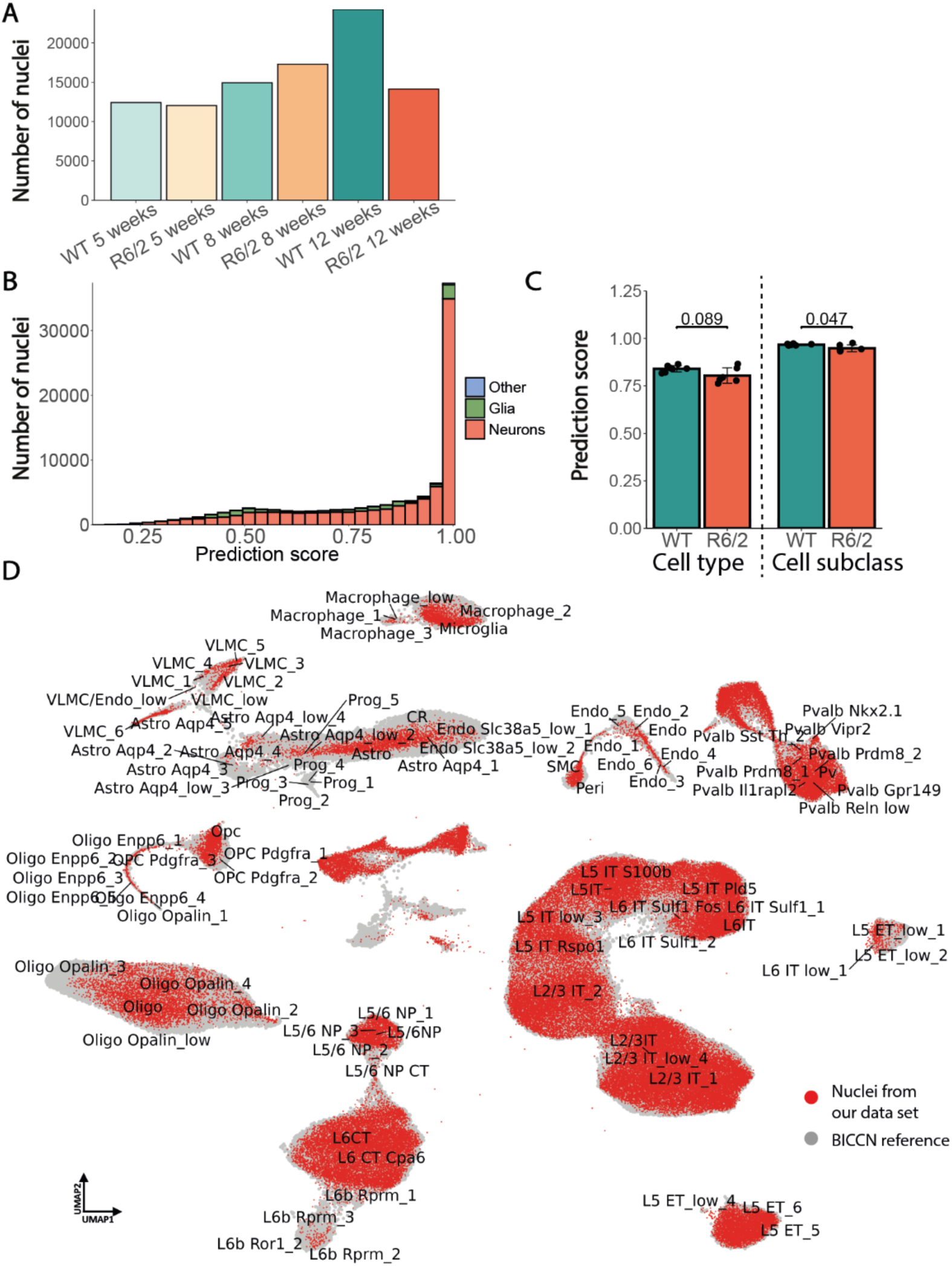
Integration of the snRNA-seq dataset into the reference atlas. **A**, Plot of nuclei numbers that passed quality control per age and genotype. **B**, Histogram of prediction scores for nuclei integrated into the reference atlas by cell type. **C**, Prediction scores for nuclei being assigned to the correct cell type and cell subclass, separated by genotype. Data are presented as mean ± standard deviation. Unpaired two-sided t-test. **D**, UMAP plot showing integration of our current dataset (red) into the primary motor cortex reference atlas (grey) (Yao et al., 2021).

**Supplementary Figure S2.**
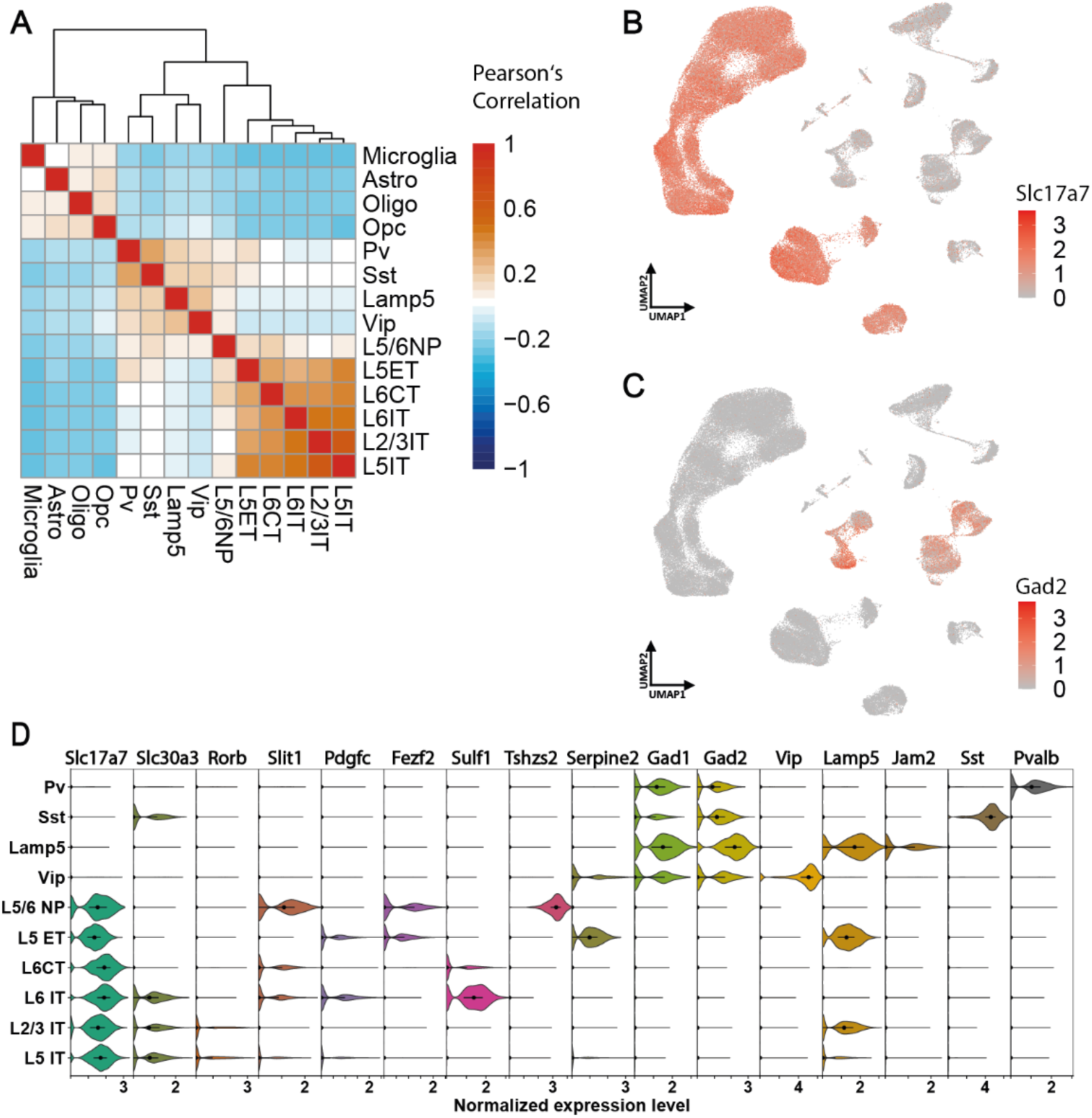
Annotation of cell subclasses in the snRNA-seq dataset. **A**, Heatmaps of pairwise Pearson’s correlation of the scaled top 4,000 variable genes between each cell subclass with hierarchical clustering by Euclidean distance. **B**, UMAP plot showing normalized expression of *Slc17a7*. **C**, UMAP plot showing normalized expression of *Gad2*. **D**, Violin plots of normalized expression of common marker genes for the indicated neuronal cell subclasses. Dots mark the median, and lines indicate the range of the second and third quartile.

**Supplementary Figure S3.**
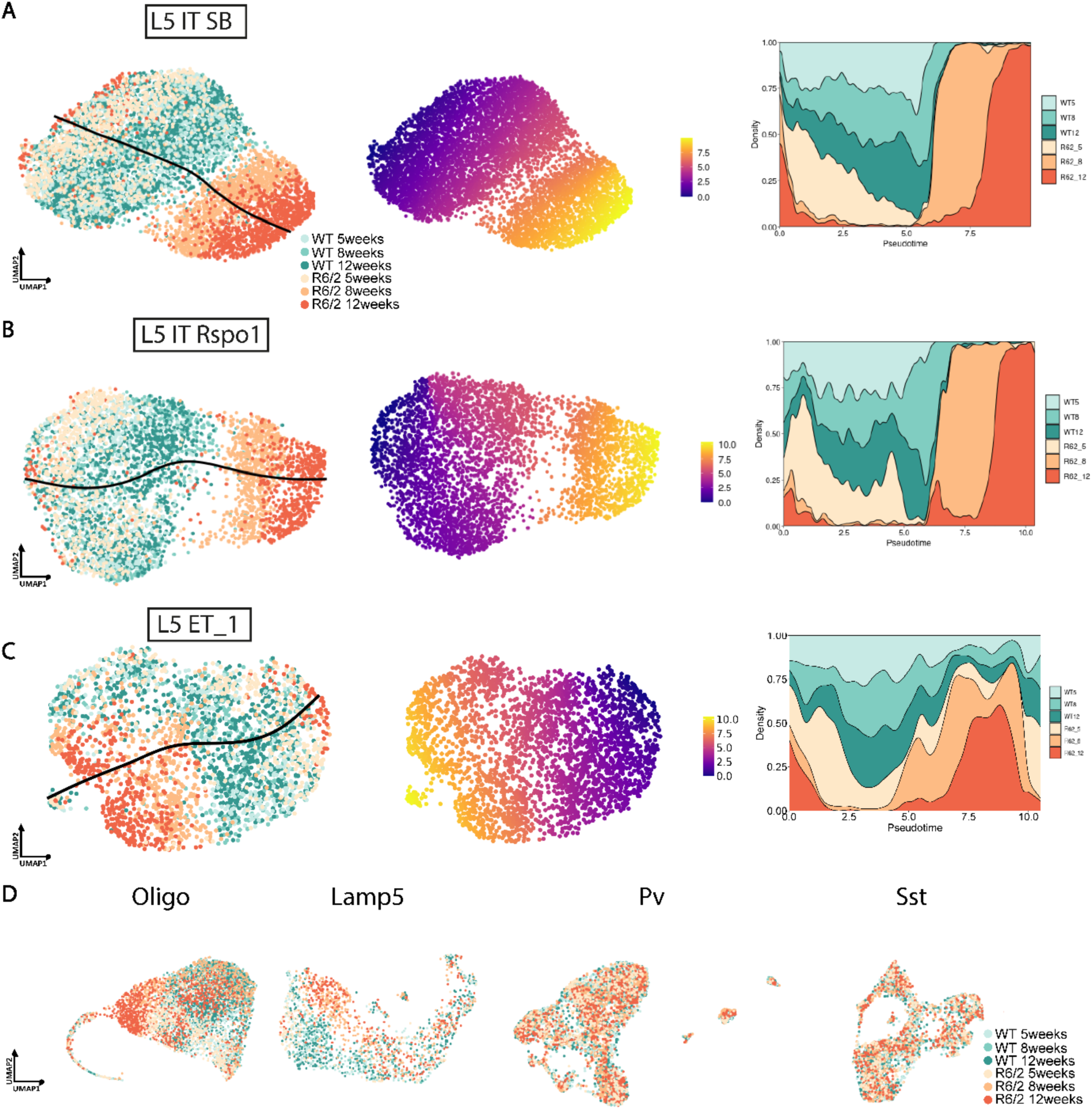
Pseudotime trajectory analysis of additional neuronal populations. **A-C**, Left: Nuclei of the L5 IT SB (top), L5 IT Rspo1 (middle), and L5 ET_1 (bottom) cell types subplotted along slingshot trajectory. Middle: Same nuclei color-coded by pseudotime value. Right: Density plots of pseudotime values showing that 5-week R6/2 nuclei from have lower pseudotime values than WT nuclei, whereas 8- and 12-week R6/2 nuclei lie at the end of the pseudotime trajectory. **D**, No clear pseudotime trajectory between R6/2 and WT nuclei of oligodendrocytes, Lamp5, Pv and Sst interneurons.

**Supplementary Figure S4.**
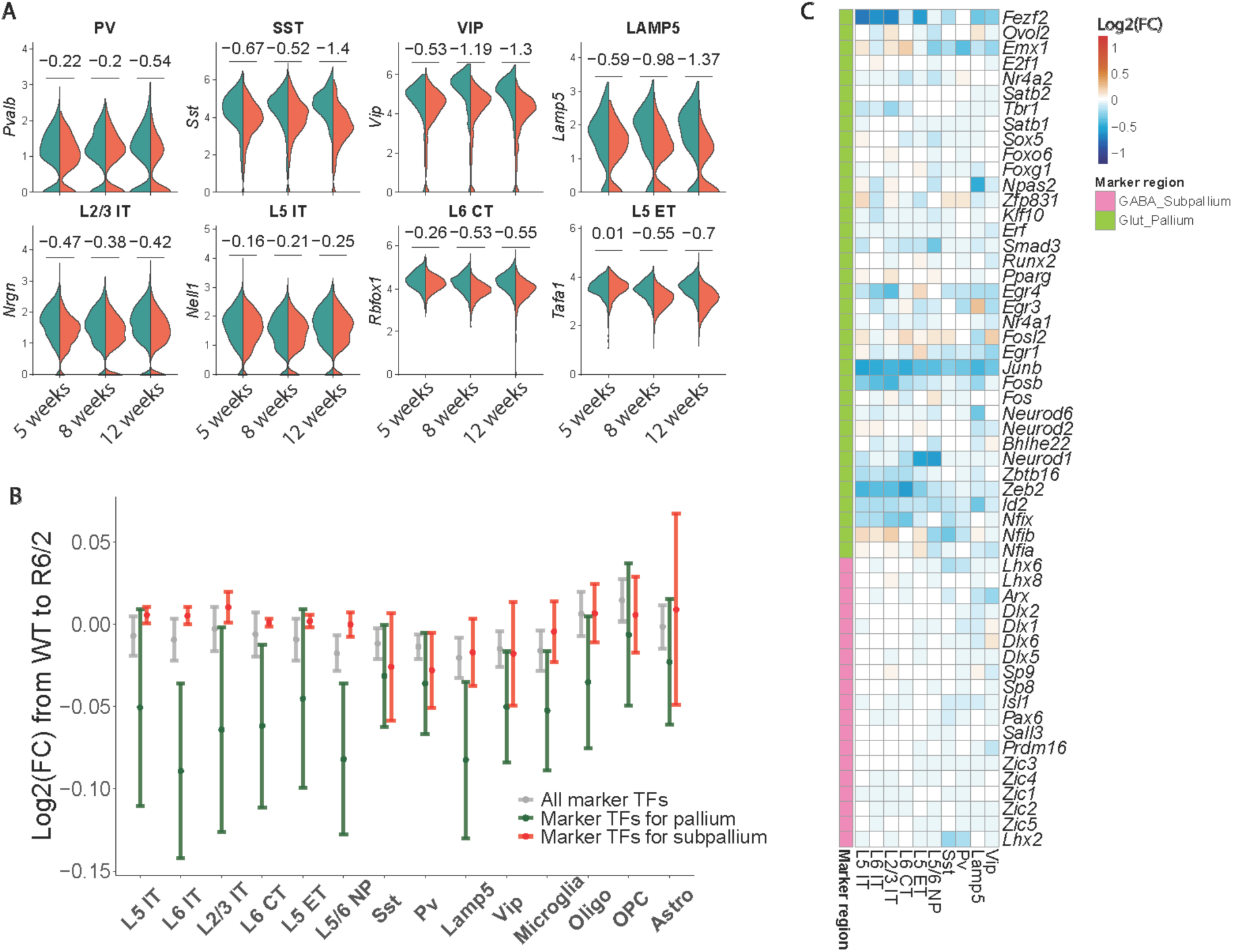
Change in expression of marker genes and transcription factors linked to cell subclass identity. **A**, Violin plots showing the expression of marker genes of the indicated cell subclasses in WT and R6/2 mice. Log2(FC) in expression from WT relative to R6/2 is indicated above the violin plot. **B**, Plot depicting the change in expression of pallium- and subpallium-specific transcription factors (Yao et al., 2023) in the indicated cell subclasses at 12 weeks of age. The dots represent the mean; error bars range the 95% confidence interval. **C,** Heat map showing the differential expression of individual transcription factors specific to the pallium and subpallium across cell subclasses at 12 weeks of age.

**Supplementary Figure S5.**
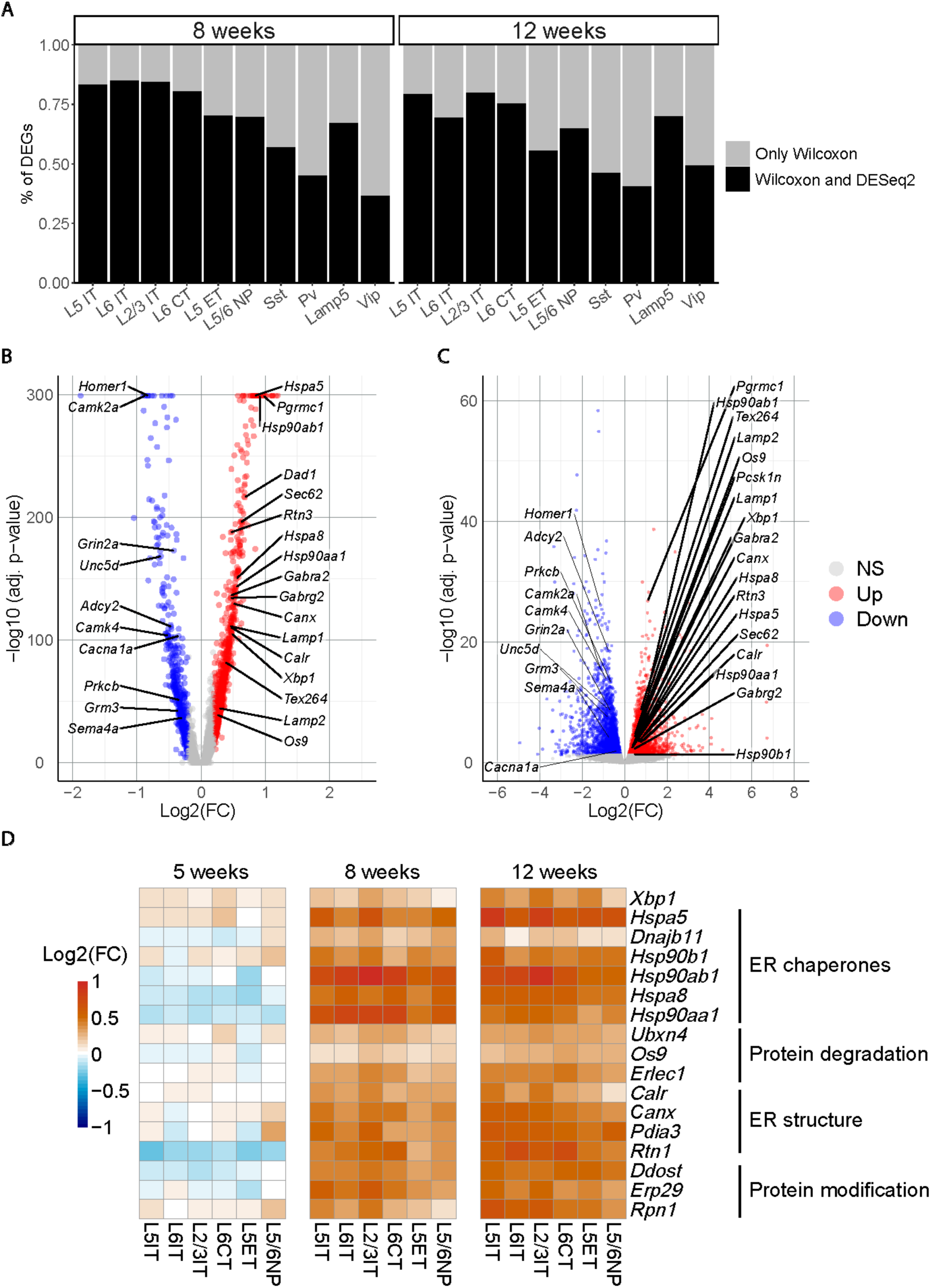
Analysis of differentially expressed genes. **A**, Percentage of DEGs identified only with Wilcoxon rank sum test and with Wilcoxon rank sum test and DeSeq2 analysis of pseudo bulk counts. Both analyses show overall consistent results. All DEGs discussed in this study were identified with both methods. **B-C**, Volcano plot depicting DEGs in L2/3 IT neurons at 12 weeks determined Wilcoxon rank sum test only (B) and by DeSeq2 of bulk counts from L2/3 IT neurons at 8 and 12 weeks combined. **D**, Heatmap showing the differential expression of ER-related genes in R6/2 mice relative to WT across all three time points.

**Supplementary Figure S6.**
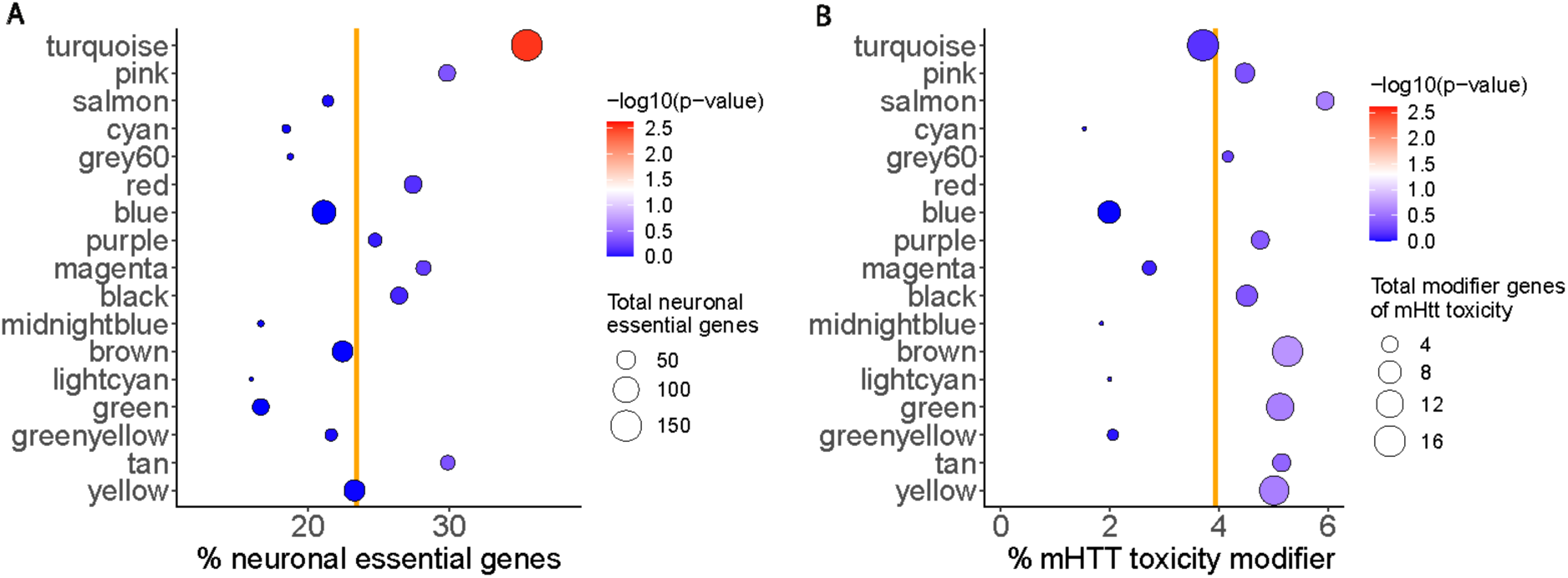
Representation of neuronal essential genes and mHTT toxicity modifiers in the modules revealed by scWGCNA analysis. Gene modules described in Fig. 4 were tested for over-representation of the 3,009 neuronal essential genes (A) and 556 mHTT toxicity modifiers (B) identified in the study by Wertz et al. (2020) and detected in our dataset. The vertical lines indicate the proportion of neuronal essential genes and mHTT toxicity modifier genes among genes expressed in neurons.

**Supplementary Figure S7.**
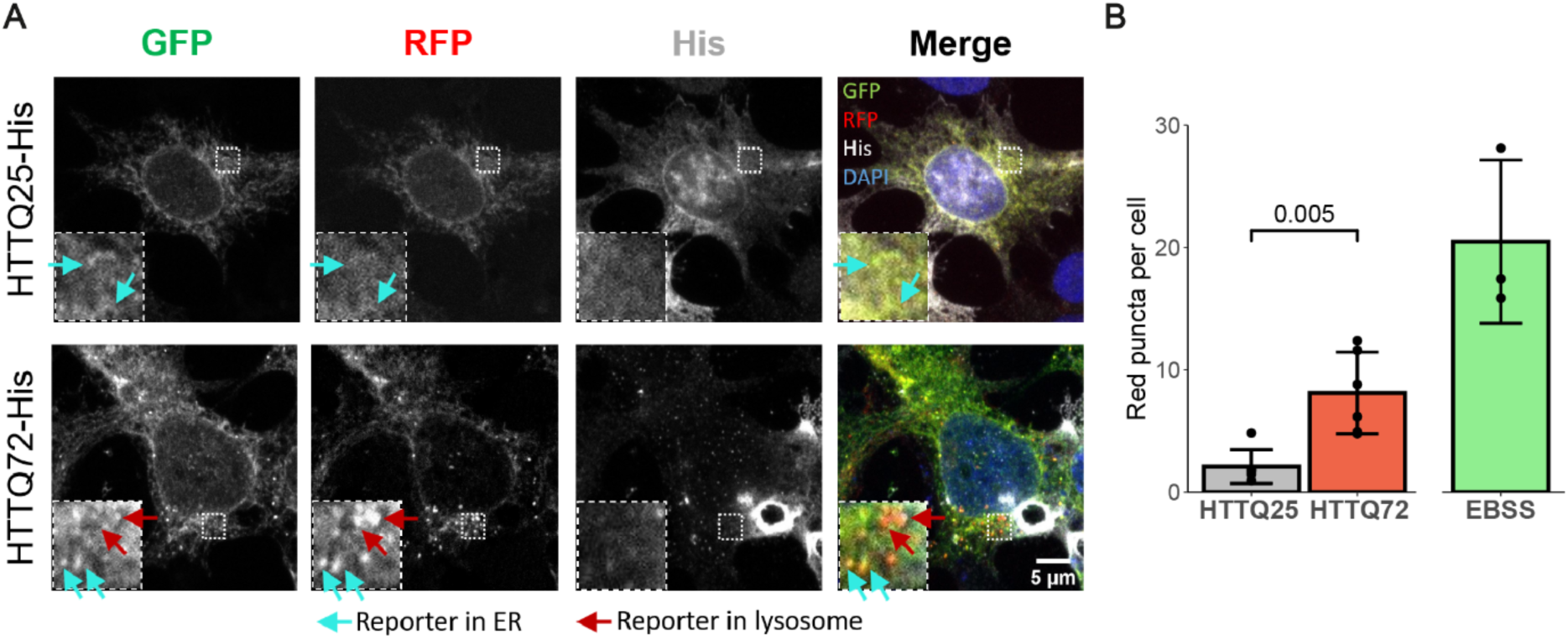
**A**, Representative images of the stable HEK cell line with the ER-phagy reporter, transfected with the indicated constructs. HTT was detected by staining against His-tag. Insets show higher magnifications of the areas marked by dashed boxes. Cyan and red arrows point to ssRFP-GFP-KDEL reporter in the ER and lysosomes, respectively. **B**, Quantificaton of red puncta corresponding to reporter inside lysosomes in the indicated conditions. Starvation of cells for 4 h in EBSS medium served as a positive control for inducing autophagy. n=7 independent experiments for HTT conditions and 3 for EBSS, 10-17 cells per replicate. Unpaired two-sided t-test.

## References

Al-Dalahmah, O., A.A. Sosunov, A. Shaik, K. Ofori, Y. Liu, J.P. Vonsattel, I. Adorjan, V. Menon, and J.E. Goldman. 2020. Single-nucleus RNA-seq identifies Huntington disease astrocyte states. Acta Neuropathol Commun. 8:19.

Barnat, M., M. Capizzi, E. Aparicio, S. Boluda, D. Wennagel, R. Kacher, R. Kassem, S. Lenoir, F. Agasse, B.Y. Braz, J.P. Liu, J. Ighil, A. Tessier, S.O. Zeitlin, C. Duyckaerts, M. Dommergues, A. Durr, and S. Humbert. 2020. Huntington’s disease alters human neurodevelopment. Science.

Braz, B.Y., D. Wennagel, L. Ratie, D.A.R. de Souza, J.C. Deloulme, E.L. Barbier, A. Buisson, F. Lante, and S. Humbert. 2022. Treating early postnatal circuit defect delays Huntington’s disease onset and pathology in mice. Science. 377:eabq5011.

Burns, M.S., R. Miramontes, J. Wu, R. Gulia, M.S. Saddala, A.L. Lau, T. Quach, J.C. Reidling, V. Swarup, A.R. La Spada, R.G. Lim, and L.M. Thompson. 2025. Distinct molecular patterns in R6/2 HD mouse brain: Insights from spatiotemporal transcriptomics. Neuron. 113:2416–2437 e2416.

Chen, Y.J., J. Knupp, A. Arunagiri, L. Haataja, P. Arvan, and B. Tsai. 2021. PGRMC1 acts as a size-selective cargo receptor to drive ER-phagic clearance of mutant prohormones. Nat Commun. 12:5991.

Chino, H., T. Hatta, T. Natsume, and N. Mizushima. 2019. Intrinsically Disordered Protein TEX264 Mediates ER-phagy. Mol Cell. 74:909–921 e906.

Chino, H., and N. Mizushima. 2020. ER-Phagy: Quality Control and Turnover of Endoplasmic Reticulum. Trends Cell Biol. 30:384–398.

Dann, E., N.C. Henderson, S.A. Teichmann, M.D. Morgan, and J.C. Marioni. 2022. Differential abundance testing on single-cell data using k-nearest neighbor graphs. Nat Biotechnol. 40:245–253.

Estrada-Sanchez, A.M., C.L. Burroughs, S. Cavaliere, S.J. Barton, S. Chen, X.W. Yang, and G.V. Rebec. 2015. Cortical efferents lacking mutant huntingtin improve striatal neuronal activity and behavior in a conditional mouse model of Huntington’s disease. J Neurosci. 35:4440–4451.

Estrada-Sanchez, A.M., and G.V. Rebec. 2013. Role of cerebral cortex in the neuropathology of Huntington’s disease. Front Neural Circuits. 7:19.

Feregrino, C., and P. Tschopp. 2022. Assessing evolutionary and developmental transcriptome dynamics in homologous cell types. Dev Dyn. 251:1472–1489.

Fernandez-Garcia, S., S. Conde-Berriozabal, E. Garcia-Garcia, C. Gort-Paniello, D. Bernal-Casas, G. Garcia-Diaz Barriga, J. Lopez-Gil, E. Munoz-Moreno, G. Soria, L. Campa, F. Artigas, M.J. Rodriguez, J. Alberch, and M. Masana. 2020. M2 cortex-dorsolateral striatum stimulation reverses motor symptoms and synaptic deficits in Huntington’s disease. Elife. 9.

Fu, H., A. Possenti, R. Freer, Y. Nakano, N.C. Hernandez Villegas, M. Tang, P.V.M. Cauhy, B.A. Lassus, S. Chen, S.L. Fowler, H.Y. Figueroa, E.D. Huey, G.V.W. Johnson, M. Vendruscolo, and K.E. Duff. 2019. A tau homeostasis signature is linked with the cellular and regional vulnerability of excitatory neurons to tau pathology. Nature neuroscience. 22:47–56.

Group, T.H.s.D.C.R. 1993. A novel gene containing a trinucleotide repeat that is expanded and unstable on Huntington’s disease chromosomes. . Cell. 72:971–983.

Gu, X., V.M. Andre, C. Cepeda, S.H. Li, X.J. Li, M.S. Levine, and X.W. Yang. 2007. Pathological cell-cell interactions are necessary for striatal pathogenesis in a conditional mouse model of Huntington’s disease. Mol Neurodegener. 2:8.

Hanquier, Z., J. Misra, R. Baxter, and J.L. Maiers. 2023. Stress and Liver Fibrogenesis: Understanding the Role and Regulation of Stress Response Pathways in Hepatic Stellate Cells. Am J Pathol. 193:1363–1376.

HD, i.C. 2017. Developmental alterations in Huntington’s disease neural cells and pharmacological rescue in cells and mice. Nature neuroscience. 20:648–660.

Hendershot, L.M. 2004. The ER function BiP is a master regulator of ER function. Mt Sinai J Med. 71:289–297.

Hill, M.A., A.M. Sykes, and G.D. Mellick. 2023. ER-phagy in neurodegeneration. J Neurosci Res. 101:1611–1623.

Hosp, F., S. Gutierrez-Angel, M.H. Schaefer, J. Cox, F. Meissner, M.S. Hipp, F.U. Hartl, R. Klein, I. Dudanova, and M. Mann. 2017. Spatiotemporal Proteomic Profiling of Huntington’s Disease Inclusions Reveals Widespread Loss of Protein Function. Cell Rep. 21:2291–2303.

Jeong, H., D.E. Cohen, L. Cui, A. Supinski, J.N. Savas, J.R. Mazzulli, J.R. Yates, 3rd, L. Bordone, L. Guarente, and D. Krainc. 2011. Sirt1 mediates neuroprotection from mutant huntingtin by activation of the TORC1 and CREB transcriptional pathway. Nat Med. 18:159–165.

Kim, D.Y., J.Y. Shin, J.E. Lee, H.N. Kim, S.J. Chung, H.S. Yoo, S.J. Kim, H.J. Cho, E.J. Lee, S.J. Nam, S.H. Kim, J. Jang, S.E. Lee, and P.H. Lee. 2023. A selective ER-phagy exerts neuroprotective effects via modulation of alpha-synuclein clearance in parkinsonian models. Proc Natl Acad Sci U S A. 120:e2221929120.

Langfelder, P., and S. Horvath. 2008. WGCNA: an R package for weighted correlation network analysis. BMC Bioinformatics. 9:559.

Leitman, J., F. Ulrich Hartl, and G.Z. Lederkremer. 2013. Soluble forms of polyQ-expanded huntingtin rather than large aggregates cause endoplasmic reticulum stress. Nat Commun. 4:2753.

Lim, R.G., O. Al-Dalahmah, J. Wu, M.P. Gold, J.C. Reidling, G. Tang, M. Adam, D.K. Dansu, H.J. Park, P. Casaccia, R. Miramontes, A.M. Reyes-Ortiz, A. Lau, R.A. Hickman, F. Khan, F. Paryani, A. Tang, K. Ofori, E. Miyoshi, N. Michael, N. McClure, X.E. Flowers, J.P. Vonsattel, S. Davidson, V. Menon, V. Swarup, E. Fraenkel, J.E. Goldman, and L.M. Thompson. 2022. Huntington disease oligodendrocyte maturation deficits revealed by single-nucleus RNAseq are rescued by thiamine-biotin supplementation. Nat Commun. 13:7791.

Lindholm, D., H. Wootz, and L. Korhonen. 2006. ER stress and neurodegenerative diseases. Cell Death Differ. 13:385–392.

Love, M.I., W. Huber, and S. Anders. 2014. Moderated estimation of fold change and dispersion for RNA-seq data with DESeq2. Genome Biol. 15:550.

Madisen, L., T.A. Zwingman, S.M. Sunkin, S.W. Oh, H.A. Zariwala, H. Gu, L.L. Ng, R.D. Palmiter, M.J. Hawrylycz, A.R. Jones, E.S. Lein, and H. Zeng. 2010. A robust and high-throughput Cre reporting and characterization system for the whole mouse brain. Nature neuroscience. 13:133–140.

Malaiya, S., M. Cortes-Gutierrez, B.R. Herb, S.R. Coffey, S.R.W. Legg, J.P. Cantle, C. Colantuoni, J.B. Carroll, and S.A. Ament. 2021. Single-Nucleus RNA-Seq Reveals Dysregulation of Striatal Cell Identity Due to Huntington’s Disease Mutations. J Neurosci. 41:5534–5552.

Mangiarini, L., K. Sathasivam, M. Seller, B. Cozens, A. Harper, C. Hetherington, M. Lawton, Y. Trottier, H. Lehrach, S.W. Davies, and G.P. Bates. 1996. Exon 1 of the HD gene with an expanded CAG repeat is sufficient to cause a progressive neurological phenotype in transgenic mice. Cell. 87:493–506.

Mathys, H., J. Davila-Velderrain, Z. Peng, F. Gao, S. Mohammadi, J.Z. Young, M. Menon, L. He, F. Abdurrob, X. Jiang, A.J. Martorell, R.M. Ransohoff, B.P. Hafler, D.A. Bennett, M. Kellis, and L.H. Tsai. 2019. Single-cell transcriptomic analysis of Alzheimer’s disease. Nature. 570:332–337.

Matsushima, A., S.S. Pineda, J.R. Crittenden, H. Lee, K. Galani, J. Mantero, G. Tombaugh, M. Kellis, M. Heiman, and A.M. Graybiel. 2023. Transcriptional vulnerabilities of striatal neurons in human and rodent models of Huntington’s disease. Nat Commun. 14:282.

McColgan, P., J. Joubert, S.J. Tabrizi, and G. Rees. 2020. The human motor cortex microcircuit: insights for neurodegenerative disease. Nat Rev Neurosci.

McGinnis, C.S., L.M. Murrow, and Z.J. Gartner. 2019. DoubletFinder: Doublet Detection in Single-Cell RNA Sequencing Data Using Artificial Nearest Neighbors. Cell Syst. 8:329–337 e324.

Molina-Calavita, M., M. Barnat, S. Elias, E. Aparicio, M. Piel, and S. Humbert. 2014. Mutant huntingtin affects cortical progenitor cell division and development of the mouse neocortex. J Neurosci. 34:10034–10040.

Myers, R.H., J.P. Vonsattel, P.A. Paskevich, D.K. Kiely, T.J. Stevens, L.A. Cupples, E.P. Richardson, Jr., and E.D. Bird. 1991. Decreased neuronal and increased oligodendroglial densities in Huntington’s disease caudate nucleus. J Neuropathol Exp Neurol. 50:729–742.

Orth, M., S. Schippling, S.A. Schneider, K.P. Bhatia, P. Talelli, S.J. Tabrizi, and J.C. Rothwell. 2010. Abnormal motor cortex plasticity in premanifest and very early manifest Huntington disease. J Neurol Neurosurg Psychiatry. 81:267–270.

Pineda, S.S., H. Lee, B.E. Fitzwalter, S. Mohammadi, L.J. Pregent, M.E. Gardashli, J. Mantero, E. Engelberg-Cook, M. DeJesus-Hernandez, M. van Blitterswijk, C. Pottier, R. Rademakers, B. Oskarsson, J.S. Shah, R.C. Petersen, N.R. Graff-Radford, B.F. Boeve, D.S. Knopman, K.A. Josephs, M. DeTure, M.E. Murray, D.W. Dickson, M. Heiman, V.V. Belzil, and M. Kellis. 2021. Single-cell profiling of the human primary motor cortex in ALS and FTLD. bioRxiv.

Praschberger, R., S. Kuenen, N. Schoovaerts, N. Kaempf, J. Singh, J. Janssens, J. Swerts, E. Nachman, C. Calatayud, S. Aerts, S. Poovathingal, and P. Verstreken. 2023. Neuronal identity defines alpha-synuclein and tau toxicity. Neuron. 111:1577–1590 e1511.

Reading, S.A., A.C. Dziorny, L.A. Peroutka, M. Schreiber, L.M. Gourley, V. Yallapragada, A. Rosenblatt, R.L. Margolis, J.J. Pekar, G.D. Pearlson, E. Aylward, J. Brandt, S.S. Bassett, and C.A. Ross. 2004. Functional brain changes in presymptomatic Huntington’s disease. Ann Neurol. 55:879–883.

Reggiori, F., and M. Molinari. 2022. ER-phagy: mechanisms, regulation, and diseases connected to the lysosomal clearance of the endoplasmic reticulum. Physiol Rev. 102:1393–1448.

Rosas, H.D., N.D. Hevelone, A.K. Zaleta, D.N. Greve, D.H. Salat, and B. Fischl. 2005. Regional cortical thinning in preclinical Huntington disease and its relationship to cognition. Neurology. 65:745–747.

Rosas, H.D., D.H. Salat, S.Y. Lee, A.K. Zaleta, V. Pappu, B. Fischl, D. Greve, N. Hevelone, and S.M. Hersch. 2008. Cerebral cortex and the clinical expression of Huntington’s disease: complexity and heterogeneity. Brain. 131:1057–1068.

Saudou, F., and S. Humbert. 2016. The Biology of Huntingtin. Neuron. 89:910–926.

Saunders, A., E.Z. Macosko, A. Wysoker, M. Goldman, F.M. Krienen, H. de Rivera, E. Bien, M. Baum, L. Bortolin, S. Wang, A. Goeva, J. Nemesh, N. Kamitaki, S. Brumbaugh, D. Kulp, and S.A. McCarroll. 2018. Molecular Diversity and Specializations among the Cells of the Adult Mouse Brain. Cell. 174:1015–1030 e1016.

Schippling, S., S.A. Schneider, K.P. Bhatia, A. Munchau, J.C. Rothwell, S.J. Tabrizi, and M. Orth. 2009. Abnormal motor cortex excitability in preclinical and very early Huntington’s disease. Biol Psychiatry. 65:959–965.

Smith, M.D., M.E. Harley, A.J. Kemp, J. Wills, M. Lee, M. Arends, A. von Kriegsheim, C. Behrends, and S. Wilkinson. 2018. CCPG1 Is a Non-canonical Autophagy Cargo Receptor Essential for ER-Phagy and Pancreatic ER Proteostasis. Dev Cell. 44:217–232 e211.

Street, K., D. Risso, R.B. Fletcher, D. Das, J. Ngai, N. Yosef, E. Purdom, and S. Dudoit. 2018. Slingshot: cell lineage and pseudotime inference for single-cell transcriptomics. BMC Genomics. 19:477.

Tabrizi, S.J., M.D. Flower, C.A. Ross, and E.J. Wild. 2020. Huntington disease: new insights into molecular pathogenesis and therapeutic opportunities. Nat Rev Neurol.

Taniguchi, H., M. He, P. Wu, S. Kim, R. Paik, K. Sugino, D. Kvitsiani, Y. Fu, J. Lu, Y. Lin, G. Miyoshi, Y. Shima, G. Fishell, S.B. Nelson, and Z.J. Huang. 2011. A resource of Cre driver lines for genetic targeting of GABAergic neurons in cerebral cortex. Neuron. 71:995–1013.

Veldman, M.B., and X.W. Yang. 2017. Molecular insights into cortico-striatal miscommunications in Huntington’s disease. Curr Opin Neurobiol. 48:79–89.

Vidal, R.L., A. Figueroa, F.A. Court, P. Thielen, C. Molina, C. Wirth, B. Caballero, R. Kiffin, J. Segura-Aguilar, A.M. Cuervo, L.H. Glimcher, and C. Hetz. 2012. Targeting the UPR transcription factor XBP1 protects against Huntington’s disease through the regulation of FoxO1 and autophagy. Hum Mol Genet. 21:2245–2262.

Voelkl, K., S. Gutierrez-Angel, S. Keeling, S. Koyuncu, M. da Silva Padilha, D. Feigenbutz, T. Arzberger, D. Vilchez, R. Klein, and I. Dudanova. 2023. Neuroprotective effects of hepatoma-derived growth factor in models of Huntington’s disease. Life Sci Alliance. 6.

Voelkl, K., E.K. Schulz-Trieglaff, R. Klein, and I. Dudanova. 2022. Distinct histological alterations of cortical interneuron types in mouse models of Huntington’s disease. Front Neurosci. 16:1022251.

Waldvogel, H.J., E.H. Kim, D.C. Thu, L.J. Tippett, and R.L. Faull. 2012. New Perspectives on the Neuropathology in Huntington’s Disease in the Human Brain and its Relation to Symptom Variation. J Huntingtons Dis. 1:143–153.

Waldvogel, H.J., E.H. Kim, L.J. Tippett, J.P. Vonsattel, and R.L. Faull. 2015. The Neuropathology of Huntington’s Disease. Curr Top Behav Neurosci. 22:33–80.

Wang, Z., J. Huang, S.P. Yang, and D.F. Weaver. 2022. Anti-Inflammatory Anthranilate Analogue Enhances Autophagy through mTOR and Promotes ER-Turnover through TEX264 during Alzheimer-Associated Neuroinflammation. ACS Chem Neurosci. 13:406–422.

Wertz, M.H., M.R. Mitchem, S.S. Pineda, L.J. Hachigian, H. Lee, V. Lau, A. Powers, R. Kulicke, G.K. Madan, M. Colic, M. Therrien, A. Vernon, V.F. Beja-Glasser, M. Hegde, F. Gao, M. Kellis, T. Hart, J.G. Doench, and M. Heiman. 2020. Genome-wide In Vivo CNS Screening Identifies Genes that Modify CNS Neuronal Survival and mHTT Toxicity. Neuron. 106:76–89 e78.

Wolf, D., C. Roder, M. Sendtner, and P. Luningschror. 2024. An Essential Role for Calnexin in ER-Phagy and the Unfolded Protein Response. Cells. 13.

Wu, T., E. Hu, S. Xu, M. Chen, P. Guo, Z. Dai, T. Feng, L. Zhou, W. Tang, L. Zhan, X. Fu, S. Liu, X. Bo, and G. Yu. 2021. clusterProfiler 4.0: A universal enrichment tool for interpreting omics data. Innovation (Camb). 2:100141.

Yang, M., S. Luo, X. Wang, C. Li, J. Yang, X. Zhu, L. Xiao, and L. Sun. 2021. ER-Phagy: A New Regulator of ER Homeostasis. Front Cell Dev Biol. 9:684526.

Yao, Z., H. Liu, F. Xie, S. Fischer, R.S. Adkins, A.I. Aldridge, S.A. Ament, A. Bartlett, M.M. Behrens, K. Van den Berge, D. Bertagnolli, H.R. de Bezieux, T. Biancalani, A.S. Booeshaghi, H.C. Bravo, T. Casper, C. Colantuoni, J. Crabtree, H. Creasy, K. Crichton, M. Crow, N. Dee, E.L. Dougherty, W.I. Doyle, S. Dudoit, R. Fang, V. Felix, O. Fong, M. Giglio, J. Goldy, M. Hawrylycz, B.R. Herb, R. Hertzano, X. Hou, Q. Hu, J. Kancherla, M. Kroll, K. Lathia, Y.E. Li, J.D. Lucero, C. Luo, A. Mahurkar, D. McMillen, N.M. Nadaf, J.R. Nery, T.N. Nguyen, S.Y. Niu, V. Ntranos, J. Orvis, J.K. Osteen, T. Pham, A. Pinto-Duarte, O. Poirion, S. Preissl, E. Purdom, C. Rimorin, D. Risso, A.C. Rivkin, K. Smith, K. Street, J. Sulc, V. Svensson, M. Tieu, A. Torkelson, H. Tung, E.D. Vaishnav, C.R. Vanderburg, C. van Velthoven, X. Wang, O.R. White, Z.J. Huang, P.V. Kharchenko, L. Pachter, J. Ngai, A. Regev, B. Tasic, J.D. Welch, J. Gillis, E.Z. Macosko, B. Ren, J.R. Ecker, H. Zeng, and E.A. Mukamel. 2021. A transcriptomic and epigenomic cell atlas of the mouse primary motor cortex. Nature. 598:103–110.

Yao, Z., C.T.J.v. Velthoven, M. Kunst, M. Zhang, D. McMillen, C. Lee, W. Jung, J. Goldy, A. Abdelhak, P. Baker, E. Barkan, D. Bertagnolli, J. Campos, D. Carey, T. Casper, A.B. Chakka, R. Chakrabarty, S. Chavan, M. Chen, M. Clark, J. Close, K. Crichton, S. Daniel, T. Dolbeare, L. Ellingwood, J. Gee, A. Glandon, J. Gloe, J. Gould, J. Gray, N. Guilford, J. Guzman, D. Hirschstein, W. Ho, K. Jin, M. Kroll, K. Lathia, A. Leon, B. Long, Z. Maltzer, N. Martin, R. McCue, E. Meyerdierks, T.N. Nguyen, T. Pham, C. Rimorin, A. Ruiz, N. Shapovalova, C. Slaughterbeck, J. Sulc, M. Tieu, A. Torkelson, H. Tung, N.V. Cuevas, K. Wadhwani, K. Ward, B. Levi, C. Farrell, C.L. Thompson, S. Mufti, C.M. Pagan, L. Kruse, N. Dee, S.M. Sunkin, L. Esposito, M.J. Hawrylycz, J. Waters, L. Ng, K.A. Smith, B. Tasic, X. Zhuang, and H. Zeng. 2023. A high-resolution transcriptomic and spatial atlas of cell types in the whole mouse brain. bioRxiv:2023.2003.2006.531121.

Zhang, F., L. Cong, S. Lodato, S. Kosuri, G.M. Church, and P. Arlotta. 2011. Efficient construction of sequence-specific TAL effectors for modulating mammalian transcription. Nat Biotechnol. 29:149–153.

